# “The Metagenomics Days”: a simplified workshop on amplicon sequencing analysis with open cloud bioinformatics for eDNA and Microbiomes

**DOI:** 10.1101/2024.12.25.630314

**Authors:** Hadj Ahmed Belaouni, Michael Stevenson, Stewart Rosell, Andrew McClure

## Abstract

The “Metagenomics Days” event was organized to enhance understanding of metagenomics and microbiome analysis among participants new to the field. This paper presents an evaluation of the course’s impact through a comprehensive survey administered before and after the event. We assessed participants’ prior knowledge, experience with bioinformatics tools, and confidence levels regarding key concepts in microbiome analysis. Word clouds generated from open-ended survey responses provided additional insights into participants’ interests and pre-course familiarity with metagenomic tools and concepts. Surveys results showed substantial improvements in participants’ confidence, particularly in building bioinformatics pipelines (+41%), understanding diversity metrics (+44.1%), and applying microbiome analysis concepts (+34.8%). Similarly, understanding of core topics like cloud computing (+28%), bioinformatics workflows (+34%), and 16S rRNA gene variable regions (+27.5%) increased significantly. The course’s impact on knowledge retention was also evaluated, with participants achieving an average of 64.87% correct responses, with 25.76% unsure answers and only 9.35% incorrect responses, highlighting the effectiveness of the course in enhancing comprehension. Overall, the survey results indicate a significant increase in participants’ confidence and knowledge acquisition, particularly in the areas of cloud computing, diversity metrics, and bioinformatics pipelines. These improvements in confidence and knowledge acquisition underscore the effectiveness of the course in bridging knowledge gaps and preparing participants for future research in the complex and rapidly evolving fields of metagenomics and microbiome analysis.

## Introduction

The “Metagenomics Days” event, held from February 19^th^ to 22^nd^, 2024, was hosted by the Agri-Food and Biosciences Institute (AFBI) in Northern Ireland. Renowned for its innovative research and teaching expertise, AFBI plays a pivotal role in advancing environmental DNA (eDNA) projects and genomic technologies, making it an ideal venue for this workshop.

The course built upon rich experiences from previous training events, notably the 2019 EMBO practical course held at the Edmund Mach Foundation in San Michele all ‘Adige, Italy, a two-week event featured an exceptional lineup of bioinformatics experts, offering comprehensive insights into genomic analyses. Additionally, the “Metagenomics Days” drew inspiration from a series of Algerian workshops organized by the Laboratoire de Biologie des Systèmes Microbiens (LBSM) at the Ecole Normale Supérieure de Kouba, Algiers, Algeria. Held between 2021 and 2023, these workshops focused on microbial phylogenetics, genomics, comparative genomics, and phylogenomics, providing a foundation for the design of this event.

This four-day event introduced participants to the core concepts and practical applications of amplicon DNA sequencing, microbiome analysis, and cloud-based bioinformatics. The course centered on a user-friendly adaptation of the “Atacama soil microbiome” tutorial (https://docs.qiime2.org/2024.10/tutorials/atacama-soils/), utilizing QIIME2’s state-of-the-art pipeline to teach amplicon sequencing analysis. A blend of lectures and hands-on sessions ensured participants gained both theoretical understanding and practical skills. Tools like QIIME2[1] and the ‘Galaxy’ cloud bioinformatics portal [2] were integral to the training, offering participants robust, cloud-based solutions for metagenomic data analysis.

The “Metagenomics Days” event was crafted with a clear objective in mind: to introduce new users to the complex yet rewarding world of metagenomics amplicon sequencing through an enjoyable and effective learning process. Tailored for participants of varying expertise levels—from beginners to seasoned researchers—the workshop provided a balanced mix of foundational knowledge and advanced techniques. The flexible organization and participant engagement made the learning experience impactful and enjoyable. By fostering an engaging and inclusive learning environment, the event ensured attendees left equipped to apply cutting-edge metagenomic methods in their own research.

This paper aims to evaluate the effectiveness of the “Metagenomics Days” event in bridging knowledge gaps and enhancing participants’ understanding of metagenomics, microbiome analysis, and bioinformatics tools, while also serving as an inspiration for institutions seeking to replicate this impactful learning experience to empower future researchers in this rapidly advancing field. Based on a series of anonymous surveys fulfilled by the participants, the responses were analyzed using a combination of descriptive and inferential statistical methods to assess pre- and post-workshop knowledge and identify significant changes in confidence and understanding levels.

## Material and methods

### 1. Communication

#### 1.1. Announcements and correspondences

Announcements and correspondences were handled through multiple channels to cater to the needs of both AFBI members and external participants, facilitating clear communication and content sharing. The primary channel for official communications with participants, lecturers, and support staff was our AFBI mailing service. This channel was used for general announcements, including registration confirmation, schedule details, and logistical information about the event.

#### 1.2. Ad Hoc workshop coordination

During the workshop, real-time coordination and content sharing were necessary to adapt to ongoing needs and challenges. To facilitate this, our personal email mailboxes were used for ad hoc communication, including last-minute changes to the schedule, sharing of additional resources, and answering participant queries.

Dr Stevenson’s professional email address served as a dedicated channel for distributing pre-course materials specifically to AFBI members. These materials included preparatory reading, software installation guidelines, and datasets needed for the hands-on exercises.

For external participants, the pre-course materials were distributed via Dr Belaouni’s personal email address. This approach helped maintain a streamlined flow of information for non-AFBI members, who required additional guidance and support to prepare for the event.

#### 1.3. Opening word, closing speech and wrap-up

The event was formally initiated with an opening speech to welcome all participants, both in-person and remote. The opening remarks highlighted the goals and objectives of the Metagenomics Days, emphasizing the importance of bioinformatics in modern microbial ecology and metagenomic studies. This speech set the tone for the event, encouraging participants to engage actively, collaborate, and take full advantage of the unique learning opportunities provided by the hands-on sessions. At the conclusion of the event, a wrap-up speech was delivered to summarize the key takeaways and achievements from the Metagenomics Days. The event concluded with a group photo session, capturing the collaborative and interactive spirit of the course, and providing a lasting memento for the participants.

### 2. Implementation strategy

#### 2.1. The registration and selection processes

The participant selection process for the “Metagenomics Days” event was designed to ensure that the attendees would benefit maximally from the program while contributing to an environment of active engagement and collaboration. A targeted approach was combined with an open registration process, maximizing both inclusivity and relevance to the event’s core focus.

In order to tailor the training content to our participants, and to gain insight about the composition of our audience, the pre-course registration form covered various aspects (supplementary material S. 1), designed as a basis for the selection process, but also to know the different expectations and backgrounds of the participants. The registration process was initiated through forms distributed across various platforms. These forms required applicants to provide information about their current academic or research status, prior experience in metagenomics and bioinformatics, and ongoing or upcoming projects related to eDNA, microbiomes, or metagenomics. This allowed the organizing committee to assess the suitability of applicants based on their background and alignment with the course content.

To maintain focus and provide a high-quality, hands-on learning experience, priority was given to individuals actively working on microbiome and metagenomics projects. The aim was to equip participants with tools they could directly apply to their work, encouraging them to bring their own datasets to the course. Preference was also given to PhD students and researchers from collaborating institutions, ensuring they would benefit from the structured, practical bioinformatics sessions provided.

In addition to open registration, targeted invitations were sent out to specific researchers and students within the “Laboratoire de Biologie des Systemes Microbiens” (LBSM lab), AFBI, and Queen’s University Belfast, fostering collaboration between institutions. Priority was given to those already involved in relevant projects, particularly in eDNA and microbiome research, ensuring that participants were well-prepared to engage with the advanced bioinformatics and sequencing topics covered during the event. This strategy ensured that the event attracted a balanced mix of early-career researchers and seasoned professionals.

This targeted yet inclusive approach helped ensure that the event was accessible to those who would benefit the most while maintaining diversity in participants’ academic and research backgrounds.

#### 2.2. The program: a structured learning environment

The “Metagenomics Days” course was structured (supplementary material S. 2) to provide participants with a comprehensive and interactive learning experience that blended theoretical knowledge with practical skills. The dual approach of integrating lectures with hands-on sessions facilitated an immersive educational environment, essential for grasping the complex concepts underlying metagenomics and amplicon sequencing.

The course was carefully designed to start with foundational lectures, which introduced participants to the principles of metagenomics and the bioinformatics tools that would be employed throughout the training. These lectures laid the groundwork for subsequent hands-on sessions, ensuring that participants had a solid understanding of key concepts before they engaged in hands-on activities (supplementary material S. 2).

On Day 1, the program commenced with participant registration and an introductory lecture on DNA sequencing technologies, microbiomes, and environmental DNA (eDNA), delivered by Dr. Hadj Ahmed Belaouni. This session provided participants with foundational knowledge on the sequencing platforms and their application in microbiome studies. In the afternoon, participants engaged in their first hands-on session on understanding sequencing outputs, led by Dr. Belaouni and Dr. Stevenson. The session focused on interpreting raw data from sequencing machines, providing participants with insights into sequencing quality and troubleshooting common errors.

Day 2 was dedicated to exploring cloud-based bioinformatics platforms. In the morning, Dr. Belaouni delivered a lecture on cloud metagenomics, highlighting the advantages and challenges of using integrated bioinformatics platforms for analyzing large-scale metagenomic data. The participants were introduced to Galaxy, a user-friendly cloud platform. The hands-on sessions in the afternoon were focused on navigating these cloud bioinformatics resources, with emphasis on practical tasks such as accessing cloud resources, running basic bioinformatics workflows, and interpreting output data.

On Day 3, the focus shifted to amplicon sequencing analysis, one of the core methods in microbiome studies[3]. Dr. Belaouni led a detailed lecture on the metagenomics amplicon sequencing pipeline, providing participants with an exemplar workflow to process 16S rRNA gene sequencing data. The afternoon session was fully practical, with participants actively using a pipeline for amplicon sequencing analysis in a cloud-based environment, using QIIME2. Dr. Belaouni and Dr. Stevenson guided participants through this complex process, ensuring that participants understood the sequencing steps from raw data to taxonomic classification.

Day 4 centered on microbiome metrics and interpreting the results of metagenomics studies. The day started with a lecture on microbiome metrics, discussing alpha and beta diversity, rarefaction curves, and other key statistical approaches. The final hands-on session provided a deep dive into QIIME2 output files, helping participants analyze and visualize their own sequencing data. The event concluded with an interactive discussion session, allowing participants to ask questions and share their insights, followed by group pictures and a formal closing by the organizers.

#### 2.3. The roles distribution

All organizers participated actively at various levels of the event (supplementary table 1). Dr. Belaouni’s main roles during the “Metagenomics Days” event included (but not limited to): Lead organizer, lecturer, and teacher. As lead organizer, it was his responsibility to decide on a multitude of points; this included the title of the event, the length of the program, the overall structure of each day, and the content which would be taught. He delegated key responsibilities to the rest of the team, which allowed him to focus primarily on creating and delivering the course content. His secondary role was that of a lecturer; each day would start with a morning lecture. During these lectures, Dr. Belaouni would allow for interactions from the attendees; however the majority of questions were saved for the end, so that the flow of the lecture was not interrupted. The other role which Dr. Belaouni took on was as a teacher. In the afternoon, hands-on sessions were carried out for all attendees. It was during these sessions where Dr. Belaouni was able to guide pairs of attendees through the examples which had been prepared in advance.

Dr Stevenson’s primary role was that of a ‘Deputy Organizer’. As Dr. Belaouni had a wide range of tasks to complete prior to and during the “Metagenomics Days” event, Dr. Stevenson took on this role so that no one person was overwhelmed. Tasks included the organization of badges for attendees, ordering of banners to be placed in key locations, creation of lecture materials, and acquisition of hospitality with specific dietary requirements. In addition to these main roles, Dr. Stevenson was also a lecturer during the course, specializing in EDGE (bioinformatics tutorial). This material was delivered to attendees, so that they would have a broad understanding of how to use this web-based version of QIIME2, in order to carry out analyses of their datasets and view associated outputs. Dr. Stevenson’s served as a demonstrator as well during hands-on sessions.

Mr. Russell’s role during the “Metagenomics Days” event was primarily as an organizer. By assisting the lead and deputy organisers, Mr. Russell was able to alleviate some of the pressure. His secondary role was in IT. This consisted of a variety of responsibilities including but not limited to; ensuring all audiovisual equipment was working correctly and synced up with the lecturers’ laptops at all times, taking questions from the online attendees to present to the lecturer post-lecture, providing supplementary materials to all attendees for the hands-on sessions.

Mr. McClure’s primary role was an organizer during the “Metagenomics Days” event, which consisted of a variety of responsibilities; including liaising with the hospitality provider, ensuring all materials for the hands-on session were available for attendees, replacing the daily banner describing the course content, and organizing seating arrangements for all in-person attendees. His secondary role was in IT; this consisted of a variety of responsibilities including providing appropriate proprietary cables for all critical hardware, setting up audiovisual software for the microphone used in the conference room, and assisting with online requests from remote attendees via live chat system.

#### 2.4. The logistics

##### Hospitality

In order for hospitality arrangements to be made, support from the divisional head (the Sustainable Agri-Food Sciences Division ‘SAFSD’, AFBI) was required. Once the details were provided (reasons for hosting the event, number of expected participants and benefits for AFBI), support was given from the divisional head, and suitable arrangements were made. As we did not have feedback data from attendees regarding their dietary requirements, we offered both vegetarian and gluten-free options for all. For afternoon breaks, coffee/tea was provided. Delivery of food and drink was arranged prior to commencement of the event each day. This allowed refreshments to arrive before the attendees, ensuring all options were made available.

##### Visual communication (posters, banners, and badges)

Posters and banners were provided by a digital printing service within our organization (AFBI). A request was made initially, for 5 posters and 2 banners to be made less than one week before the commencement of the event. Our requests were met promptly allowing us to organize and setup the locations of each poster/banner well in advance of the event. Name badges were also arranged. Badges were handed out on the first day, allowing everyone to engage easily with one another, and identify colleagues with whom they had previously corresponded, but not yet met in person. They were also used to allow the organizers to split up attendees based on their metagenomics experience, ensuring that each group had at least one experienced member present.

#### 2.5. Breaks and discussion: flexible pauses and engaged discussions

The hands-on sessions served as the core of the learning experience, allowing participants to apply the theoretical knowledge gained during lectures. Each practical session commenced with a brief overview of the tools then the tasks at hand, reinforcing the principles discussed earlier. Participants were encouraged to engage actively in discussions during these sessions, fostering a collaborative learning atmosphere. This interaction provided an opportunity for participants to share insights related to their own research projects, allowing the organizers to tailor discussions to their specific interests and needs. We implemented flexible pauses during the course to accommodate the intensity of the various segments. Recognizing that participants had differing levels of familiarity with bioinformatics, these breaks allowed for questions and clarifications, ensuring that everyone remained engaged and understood the material.

#### 2.6. The hands-on sessions (highlights)

##### Tools descriptions

In the initial stage of each hands-on session, we provided detailed explanations of the various bioinformatics tools to be utilized, such as QIIME2, EDGE bioinformatics[4], and additional user-friendly software like MEGA X[5], FinchTV[6], and BioEdit[7]. These tools were introduced not only to familiarize participants with their functions but also to explain their significance within the broader context of metagenomic analysis. This preparatory phase was crucial in helping participants understand the workflow of amplicon sequencing analysis, including data processing and interpretation.

##### File types and metadata preparation

We covered the basic file types commonly handled in a typical amplicon sequencing analysis, ensuring that participants were well-versed in the terminology and data formats they would encounter in their work. Furthermore, preparing a metadata file is a critical step in any metagenomic analysis[8], and we dedicated time to exploring this aspect in detail. Participants were instructed on how to structure metadata files, including necessary columns and formatting, which is essential for successful analysis using tools like QIIME2. Understanding the relationship between sample information and analysis results empowers researchers to make informed decisions about their studies.

##### Overlapping reads and quality control

To facilitate a deeper understanding of the mechanics of sequence processing, we demonstrated the process of merging reads, emphasizing the importance of paired-end sequencing[9]. Using intuitive software such as Mega, FinchTV, and BioEdit, we showcased simple examples of how overlapping reads are combined. This hands-on exploration allowed participants to visualize the merging process and grasp the principles underlying this critical step in metagenomic analysis.

##### Quality control reports

Quality control (QC) is paramount in ensuring the integrity of sequencing data[10]. Throughout the course, we examined various next-generation sequencing (NGS) libraries and their QC reports using FastQC[11]. Participants learned how different libraries exhibit distinct quality profiles, highlighting that certain quality metrics might be acceptable for one type of library while posing issues for another. For example, we discussed how high duplication levels might be problematic for genomic libraries, while they may be tolerable for amplicon libraries. This nuanced understanding of QC metrics is crucial for participants to critically assess their own sequencing data and make informed decisions about data usability.

##### Data visualization

A significant portion of the course was dedicated to data visualization techniques, essential for interpreting the results of metagenomic analyses. We introduced participants to the structure and interpretation of basic and advanced charts, such as box plots and alpha rarefaction curves. Participants learned how to read these visual representations, focusing on how to set depth thresholds for even sampling and understand their implications for microbial diversity analysis.

##### Alpha rarefaction curves and box plots

The alpha rarefaction curve was emphasized as a vital tool for assessing the relationship between sequencing depth and observed diversity[12]. By guiding participants through examples, we illustrated how these curves can help determine the adequacy of sequencing efforts in capturing the full diversity of microbial communities. Additionally, box plots were introduced to convey distribution patterns in datasets, emphasizing the importance of understanding variance in microbial abundances[13].

##### The QIIME2 “Atacama soil microbiome” tutorial as the foundation of learning amplicon sequencing analysis

The cornerstone of our pedagogical approach was the adaptation of the QIIME2 Atacama exercise to the EDGE bioinformatics platform. This innovative strategy was central to our mission of making metagenomics accessible, engaging, and impactful for the course participants.QIIME2, a leading tool for microbial community analysis, has long been the gold standard for amplicon sequencing analysis[14]. The Atacama[15] tutorial provides a comprehensive introduction to the analytical pipeline for processing metagenomic data[16]. It covers all critical steps, from raw data import, sequence quality control, and operational taxonomic unit (OTU) clustering to taxonomic classification and diversity analyses. Despite its thoroughness, the QIIME2 environment—being largely command-line based—poses significant challenges to new users, especially those without prior experience in bioinformatics or computational biology. Recognizing this barrier, we sought to adapt the Atacama tutorial for use on EDGE bioinformatics, a cloud-based platform known for its user-friendly interface. EDGE offers a streamlined and visually intuitive environment that eliminates the steep learning curve associated with command-line tools like QIIME2. Importantly, this adaptation did not dilute the scientific rigor of the Atacama tutorial. Instead, it allowed us to present the same robust bioinformatic processes in a way that was more approachable for beginners, enabling them to focus on learning the biological insights rather than struggling with computational syntax.

For this course, we pre-configured the QIIME2 Atacama pipeline on EDGE, which included steps for:

- Importing raw sequencing data
- Running sequence quality control with DADA2
- Conducting taxonomic classification
- Computing alpha and beta diversity metrics
- Visualizing results through heatmaps, bar plots, and ordination diagrams

The transition to EDGE streamlined these processes, allowing participants to interact with the analytical workflow in a much more intuitive manner. By removing the need to code, users could instead focus on understanding key concepts such as OTU clustering, phylogenetic tree construction, and taxonomic assignment. This hands-on experience helped reinforce the biological meaning behind the data they were processing, which we believe is a critical element in effective bioinformatics education.

#### 2.7. Course material and resources

At the conclusion of the course, we provided participants with comprehensive course materials, including datasets, lecture slides, and handouts summarizing key concepts (see supplementary material S. 3).

### 3. Pre- and post-course surveys analysis

To be able to evaluate the confidence levels prior and after the training but also to evaluate in an objective and anonymous way the acquired knowledge throughout the lectures and the hands-on session, we opted for a large set of questions in forms, that covered various aspects (supplementary material S. 1).

#### 3.1. Surveys organization

We evaluated the effectiveness of the workshop through online (Microsoft Forms) pre-course and post-course surveys. General pre-course surveys covered all 43 participants, both remote and in place attendees. A total of 12 in-place attendees were evaluated through post-course surveys, for confidence shifts (7 Likert-scale questions) and knowledge acquisition (13 questions). Post-course surveys included a combination of Likert-scale questions (scored from 0 to 10) and knowledge-based queries, reflecting key competencies in microbiome analysis. These surveys aimed to measure participants’ confidence in understanding key concepts, their theoretical knowledge in areas such as alpha and beta diversity, diversity metrics, and 16S rRNA gene variable regions, and their ability to apply learned skills.

#### 3.2. Surveys statistical analysis

The data collected from the pre- and post-assessments were subjected to rigorous statistical analysis to determine the significance of any changes in confidence and understanding before and after the workshop. For each question, mean, standard deviation, and range of scores were calculated. Descriptive statistics were computed to summarize the overall trends in confidence and understanding. The Shapiro-Wilk test was applied to assess the distribution, and subsequently a parametric or non-parametric test was applied to assess the differences. For all applied statistic tests, results were interpreted at a significance level of 𝑝 < 0.05. The p-values were represented in the visualizations using conventional staring scale: p-value < 0.001: *** (3 stars, highly significant); 0.001 ≤ p-value < 0.01: ** (2 stars, moderately significant); 0.01 ≤ p-value < 0.05: * (1 star, significant); p-value ≥ 0.05: no stars (not significant).

##### Post-course evaluations

###### Conceptual understanding and confidence assessments

Participants were asked to rate their confidence and understanding levels both before and after the workshop, through a series of Likert-scale questions with scores ranging from 1 (no confidence/poor understanding) to 10 (complete confidence/thorough understanding). When normality condition met, to evaluate whether there was a statistically significant change in participants’ confidence or understanding before and after the workshop, a paired t-test (parametric test) was performed for each key concept, and Cohen’s d was used to calculate the effect size (interpretation: small effect < 0.2, medium effect 0.2-0.5, large effect > 0.5). If normality was violated, a permutation test (non-parametric test) was used instead, and rank-biserial correlation was used to assess the significance and effect size of changes in confidence (interpretation: small effect < 0.1, medium effect 0.1-0.3, large effect > 0.3).

###### Knowledge acquisition evaluation

To assess participants’ responses after completing the course, three categories of responses were analyzed: Correct, False, and Not Sure. These categories represent the percentage of answers provided for each question (13 questions). Since the data were paired (percentages of Correct, False, and Not Sure responses for the same questions) and non-normally distributed (from Shapiro-Wilk test), a Friedman test was initially used to determine overall differences between the three categories. For pairwise comparisons between response categories, Wilcoxon Signed-Rank tests were applied. To quantify the magnitude of differences for the Wilcoxon Signed-Rank tests, the rank-biserial correlation was calculated as an effect size measure. The rank-biserial correlation rescales the Wilcoxon statistic to a correlation-like effect size, and depending on values the effect was qualified as: small effect (𝑟 ≈ 0.1), medium effect (𝑟 ≈ 0.3), or large effect (𝑟 ≥ 0.5).

### Software, implementation and visualization

All statistical analyses were performed using Python 3.8 with the Scipy, Pandas, Matplotlib, Seaborn, and Numpy libraries for data manipulation, statistical testing and outputs plotting. Changes in confidence and understanding were visualized using pie charts, bar charts, and box plots generated in Plotly (https://chart-studio.plotly.com), which provided a clear representation of the distribution of scores and the magnitude of improvement across the various concepts assessed.

## 4. Ethical considerations

This study was conducted in compliance with ethical guidelines for educational research. Informed consent was obtained from all participants prior to their involvement in the workshop, with assurances that all data would be anonymized and used exclusively for research purposes.

## III. Results

### 1. Enhancing user experience with QIIME2 high-quality visual outputs

One of the major challenges in teaching metagenomics is helping participants navigate the large volumes of data that such analyses generate. Raw sequencing reads, OTU tables, and taxonomic classifications can be overwhelming, particularly for new users. To address this, we took full advantage of EDGE’s robust visualization capabilities. Participants could view their results as interactive bar plots, PCoA plots, and heatmaps, which made interpreting the data more intuitive. For example, instead of manually generating complex R or Python scripts to visualize microbial diversity, participants could simply select parameters from drop-down menus within EDGE and instantly see the impact of their choices.

### 2. Optimization and problem solving in bioinformatics analyses

Another key aspect of our course strategy was the emphasis on troubleshooting and iterative analysis, which we integrated into the sessions in real time. Bioinformatics is inherently prone to issues related to data quality[17], parameter selection[18], and tool compatibility[19]. Rather than shielding participants from these challenges, we encouraged them to engage with the problems directly. Using EDGE’s simplified interface, participants could tweak parameters like sequence truncation length, taxonomic databases, and rarefaction depth to observe how these changes impacted the results. This iterative approach to data analysis gave participants a deeper understanding of the importance of parameter optimization in bioinformatics workflows. This approach during hands-on sessions proved invaluable, as participants learned not just how to follow a predefined protocol but how to think critically about their data and the underlying assumptions in bioinformatics analyses.

### 3. Broadening the impact of learning

The course curriculum was designed to be flexible enough to cater to the broad range of research interests represented by the participants. From environmental microbiomes and agricultural soil health to animal microbiomes and eDNA-based biodiversity monitoring, participants came from various scientific backgrounds. As a result, we structured the explanation of exercises in such a way that users, once they understood the Atacama exercise, were fully equipped to customize their analyses according to their own datasets and research questions. For example, participants interested in microbiome research could apply QIIME2’s advanced taxonomic classification algorithms to their own microbial datasets, while those focusing on environmental DNA (eDNA) projects were able to use EDGE to identify rare taxa and explore biodiversity metrics in their samples. This flexibility ensured that each participant could derive maximum value from the course, regardless of their specific area of study.

### 4. User feedback and participant engagement

A critical aspect of the “Metagenomics Days” course was the active engagement of participants through continuous feedback loops. We encouraged users to share their thoughts, ask questions, and provide insights throughout the sessions, which enabled us to tailor our approach in real time. One of the most consistent pieces of feedback was the appreciation for the accessibility and user-friendliness of the Galaxy portal, and the EDGE bioinformatics platform, particularly when compared to traditional command-line tools like QIIME2.

### 5. Leveraging cloud resources for metagenomics analysis

A major highlight of the course was our introduction of cloud-based bioinformatics workflows, which significantly expanded the reach and practicality of the course material. Metagenomics is computationally demanding, requiring substantial processing power[20] for tasks such as quality control, sequence denoising, and sequences clustering. By using the web-based bioinformatics platforms, we were able to alleviate the need for participants to have access to local high-performance computing environments. The cloud platform allowed them to offload these resource-heavy tasks onto remote servers, democratizing access to bioinformatics for participants from diverse geographic and economic backgrounds.

### 6. Flexibility and tool Integration

A key take away from the course was the importance of flexibility in utilizing bioinformatics tools. We demonstrated how various tools can be integrated into a coherent workflow, illustrating that there is often more than one way to achieve analytical goals in metagenomics. This adaptability encourages researchers to explore multiple avenues for analysis, fostering a mindset that values resourcefulness and innovation in the field of bioinformatics.

### 7. Forms analysis and feedback

The survey and registration data provide significant insights into the participants’ backgrounds, experiences, and the impact of the “Metagenomics Days” event on their learning. The detailed analysis below is broken into key themes:

#### 7.1. Participant demographics

##### Geographical distribution

The majority of participants came from two main regions (fig. 1a): the UK (18 participants) and Algeria (25 participants). This further underscores the global relevance of microbiome research.

**Figure 1.**
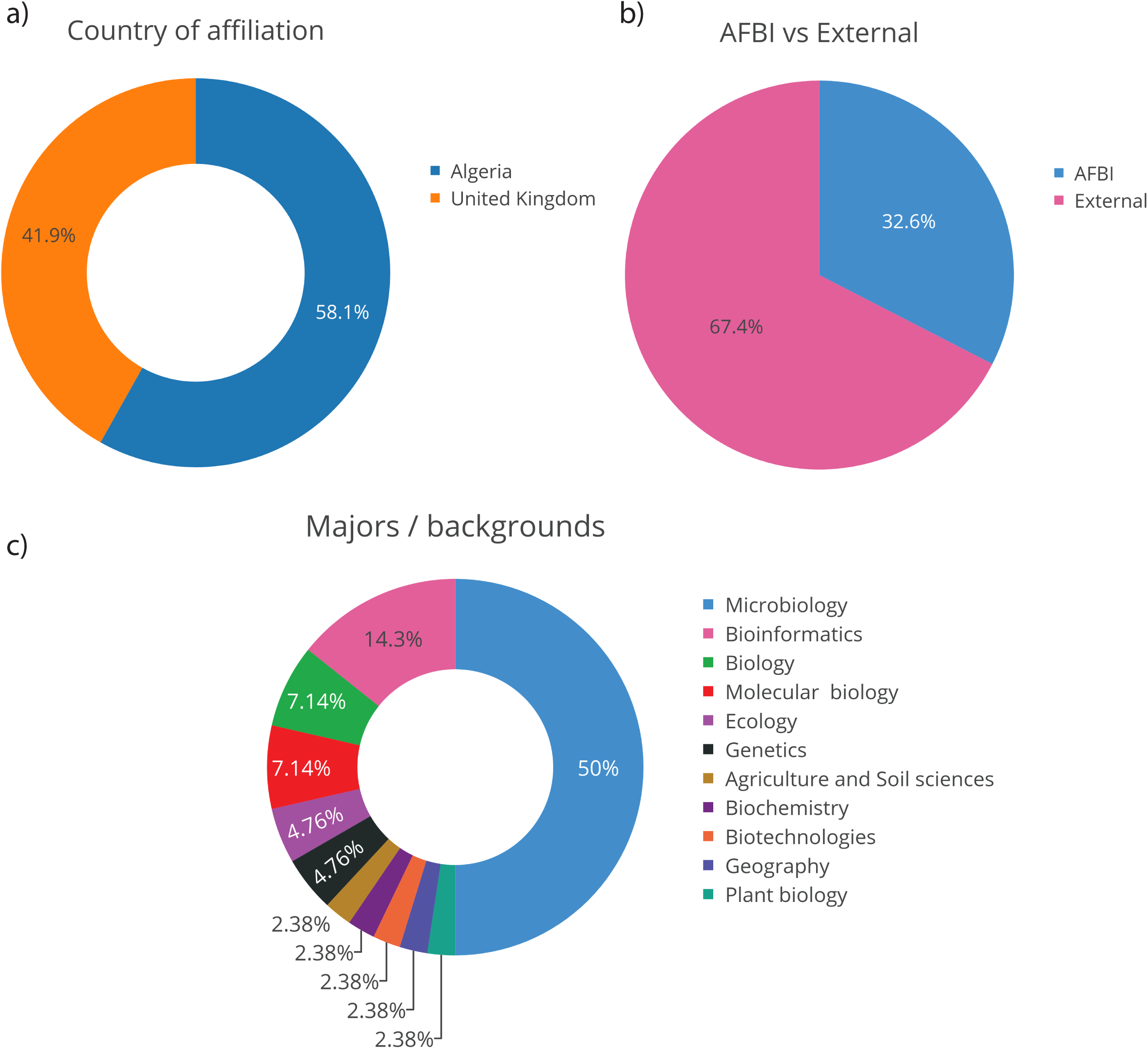
Demographic and professional background of course participants. **a)** ‘Country of affiliation’ –Pie chart showing percentage of attendees and their country of affiliation, blue refers to Algeria& orange refers to United Kingdom. **b)** ‘AFBI vs External’ - Pie chart showing percentage of attendees working internally within AFBI and those who are based externally, blue refers to AFBI & orange refers to external. **c)** ‘Majors/backgrounds’ - Pie chart showing percentage of attendees associated with specific University Degree Majors or education backgrounds.

##### Institutional and disciplinary backgrounds

A notable number of participants (14) were affiliated with AFBI, while the remaining 29 were external participants (fig. 1b). Most participants had a background in biological sciences, particularly microbiology (21 participants) (fig. 1c). This concentration reflects the workshop’s alignment with the needs of microbiologists looking to expand their expertise in bioinformatics and metagenomics.

#### 7.2. Participant experience prior to the course

##### Familiarity with OMICS and bioinformatics solutions

Nearly half (53.5%) of the attendees stated a past experience with OMICS approaches (fig. 2a), and more than half (55.8%) reported a familiarity with NGS technologies (fig. 2b); underscoring their preparedness to learn metagenomics related concepts. Approximately half of the participants had not previously used cloud-based bioinformatics solutions (fig. 2c), indicating a diverse group with varying levels of prior experience. Only approximately the third of the participants (35%) reported a past experience in using Galaxy platform for bioinformatics, despite the overwhelmingly growing popularity of the later. A similar proportion (30%) reported a certain familiarity with the use of MEGA evolutionary software, dedicated to the exploration of evolutionary relationships, and even less for the rest of the proposed tools in this question (≤ 10%) (fig. 2d). This highlights the course’s importance in introducing modern, cloud-based platforms like Galaxy and QIIME2 to those less experienced in the field. A similar split was observed for prior knowledge of microbiome data repositories/databases (fig. 2f) (5 out of 10 had no prior knowledge), reinforcing the course’s foundational role in introducing participants to relevant resources. Notably, 70% of participants were unaware of QIIME2 prior to the course (fig. 2e). This underscores the value of providing hands-on experience with specific bioinformatics tools, which many participants likely encountered for the first time during the event.

**Figure 2.**
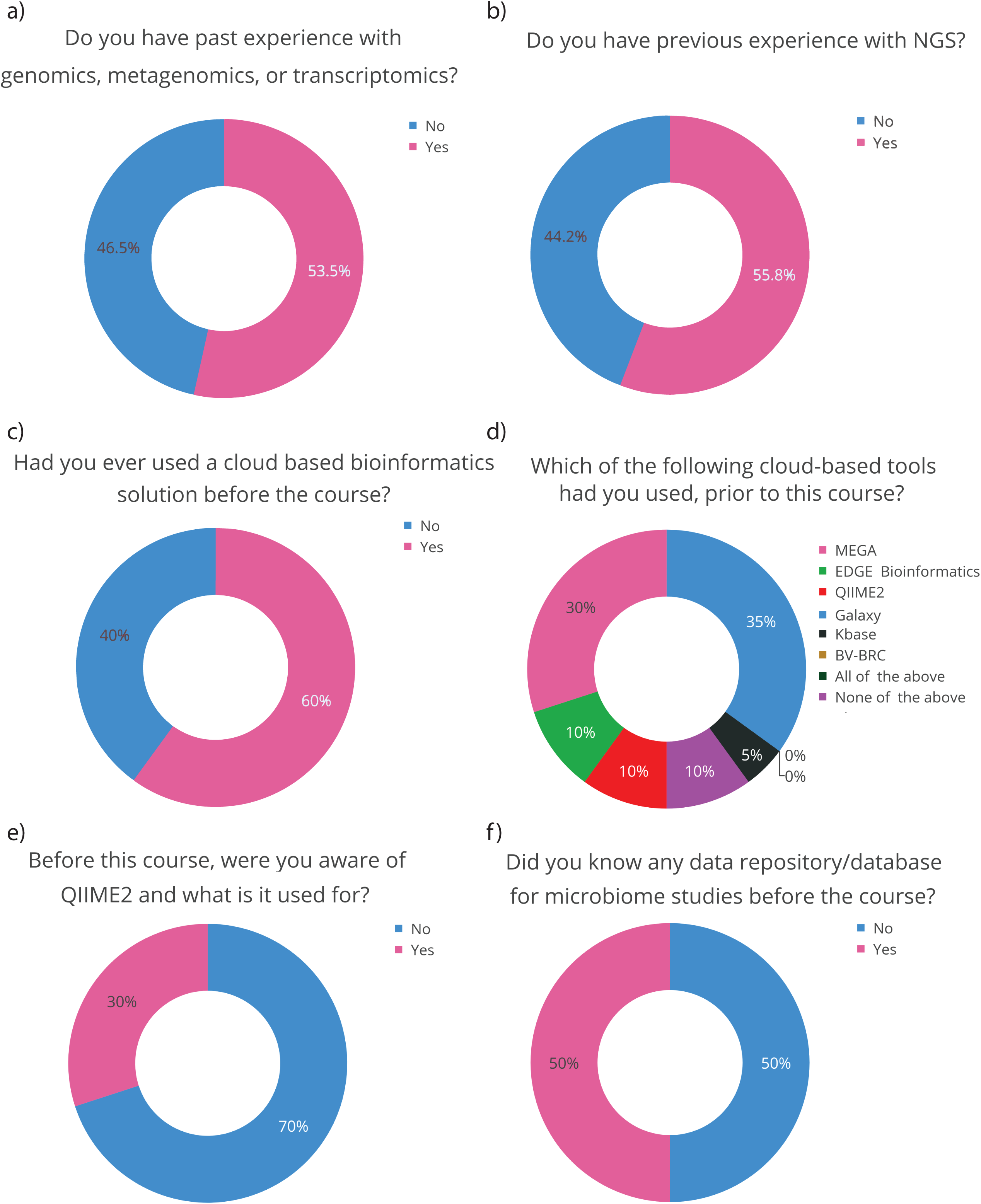
Participants’ pre-course knowledge and experience with bioinformatics tools and techniques. **a)** Pie chart showing percentage of attendees who have past experience with various -omics (genomics, metagenomics, transcriptomics), blue shows ‘no’, pink shows ‘yes’ for all charts, except Figure 2d. **b)** Pie chart showing percentage of attendees who have previous experience with Next-generation sequencing (NGS). **c)** Pie chart showing percentage of attendees who have used cloud-based bioinformatics solution prior to attending the course. **d)** Pie chart showing percentage of attendees who have used various cloud-based bioinformatics tools. **e)** Pie chart showing percentage of attendees who were aware of QIIME2 and its utility prior to attending the course. **f)** Pie chart showing percentage of attendees who were aware of any databases relating to microbiome studies prior to attending the course.

##### Bioinformatics and informatics skills

Informatics skills (fig. 3a) were moderately rated (5.25 ± 2.32, on a scale of 10), while bioinformatics-specific skills were rated lower (4.32 ± 2.25, on a scale of 10), indicating that many participants were still in the early stages of developing the specialized skill sets required for microbiome data analysis. This reinforces the need for further bioinformatics training and highlights the potential impact of continued educational efforts like the “Metagenomics Days” course. No significant difference was recorded between informatics and bioinformatics self-assessed skills (permutation test’s p-value = 1.000) (tab. 1). Participants reported an overall openness to bring their own laptop (83.7%), which facilitated the immersive experience, granted by the use of a familiar IT solution (supp. fig. 1). Moreover, most of the users reported their preference of a Windows OS (86%), compared to other systems (supp. fig. 2).

**Figure 3.**
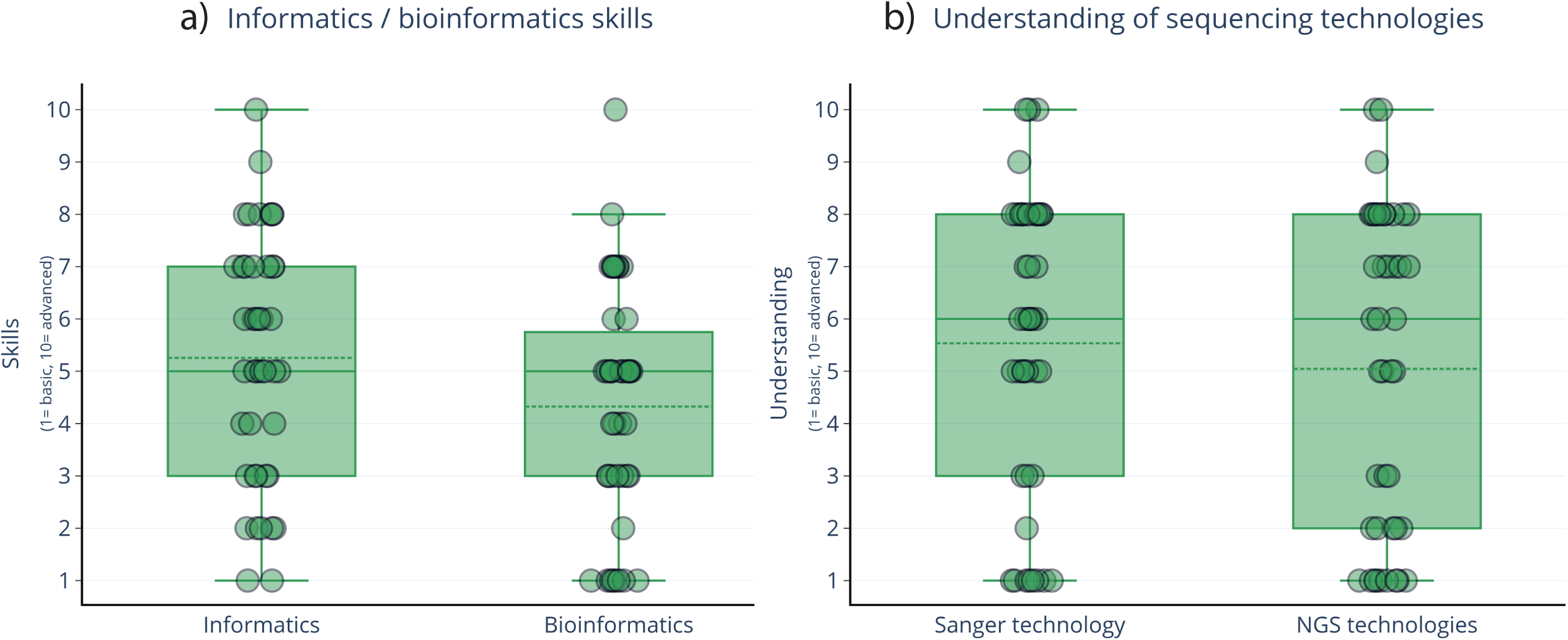
Pre-course self-evaluation of participants’ informatics and bioinformatics skills; **a)** Box plots showing self-assessment of attendees’ skills in both categories of informatics and bioinformatics. For each box: upper square line = upper quartile (q3), lower square line = lower quartile (q1), middle square line = median, upper whisker = highest observed value, lower whisker = lowest observed value; dashed middle line = mean. Skills range from 1 (basic) to 10 (advanced). **b)** Box plots showing self-assessment of attendees’ understanding of both Sanger technology and Next-generation sequencing (NGS) technologies. Understanding ranges from 1 (basic) to 10 (advanced). The confidence ratings are presented as mean ± standard deviation. A permutation test was used to compare the confidence levels in the two areas, and effect size was calculated using the rank-biserial correlation. No significant difference was observed between compared elements of the two questions.

**Table 1.** Summary of statistical analyses for participants’ confidence and knowledge acquisition before and after the course. Descriptive statistics (mean ± SD) are presented alongside test results (p-values) and effect sizes. Significant improvements were observed in post-course confidence across various categories, with large effect sizes indicating the course’s effectiveness in enhancing participants’ confidence and understanding.

| Category | Metric | Mean ± SD | | Test/<br>Statistic | P-value | Effect Size |
| --- | --- | --- | --- | --- | --- | --- |
|  |  | Before | After |  |  |  |
| **Pre-Course Surveys** | Informatics confidence | 5.25 ± 2.32 | - | Permutation Test | 1 | - |
| | Bioinformatics confidence | 4.32 ± 2.25 | - | - | - | - |
| | Sanger Technology confidence | 5.53 ± 2.86 | - | Permutation Test | 1 | - |
| | NGS Technology confidence | 5.04 ± 3.02 | - | - | - | - |
| **Post-Course Surveys** | Pipeline/Workflow confidence | 3.40 ± 2.36 | 6.80 ± 2.04 | Permutation Test | 0.047 | Rank-Biserial: 1.000 (Large) |
| | Microbiome concepts confidence | 5.16 ± 2.58 | 8.25 ± 1.60 | Permutation Test | 0.016 | Rank-Biserial: 1.000 (Large) |
| | Metadata file preparation | 2.83 ± 2.08 | 6.91 ± 1.92 | Permutation Test | 0.016 | Rank-Biserial: 1.000 (Large) |
| | Diversity metrics confidence | 3.00 ± 2.59 | 7.41 ± 2.42 | Permutation Test | 0.016 | Rank-Biserial: 1.000 (Large) |
| | Cloud computing confidence | 4.80 ± 2.97 | 7.60 ± 1.34 | Paired t-Test (t=-4.023) | 0.003 | Cohen's d: -1.272 (Large) |
| | Alpha/Beta diversity confidence | 3.25 ± 2.63 | 7.08 ± 3.05 | Permutation Test | 0.016 | Rank-Biserial: 1.000 (Large) |
| | 16S rRNA variable regions | 5.08 ± 2.46 | 7.83 ± 1.99 | Permutation Test | 0.027 | Rank-Biserial: 1.000 (Large) |
| **Overall** | Average confidence shift | 3.93 ± 1.03 | 7.41 ± 0.52 | Paired t-Test (t=-14.217) | 0 | Cohen's d: -5.374 (Very Large) |
| | Knowledge Acquisition (Quiz Scores) | 64.87 ± 23.03% (Correct) | - | Friedman Test ($\chi^2=16.000$) | 0 | Correct vs. False: r =0.956 (Large),<br>Correct vs. Unsure: r =0.802 (Large)<br>False vs. Not Sure: r = 0.967 |

##### Understanding of sequencing technologies

Participants had a moderate understanding of Sanger sequencing technology (5.53 ± 2.86, on a scale of 10) and a slightly lower understanding of NGS technologies (5.04 ± 3.02, on a scale of 10) before the course (fig. 3b). The results showed no statistically significant difference between the two technologies’ ratings before the workshop (permutation test’s p value = 1.000) (tab. 1). These values suggest that while many participants had some basic knowledge, there was still room for growth, particularly in relation to modern sequencing methods such as NGS. A large number of participants (19 out of 43) had prior experience with NGS (fig. 2b), but the data suggest that this was not necessarily accompanied by deep bioinformatics skills, given the lower self-reported confidence in building pipelines (2.7 ± 2.00, on a scale of 10) (supp. fig. 3) prior to the Galaxy tutorial.

#### 7.3. Participants’ current interests and objectives

##### Participant’s current interests

The survey conducted for the advanced bioinformatics and metagenomics course revealed a broad range of research interests (fig. 4), reflecting trends in microbial genomics, eDNA, and related fields. Many participants are engaged in eDNA projects, highlighting its growing role in environmental research and species detection, aligning well with the course’s focus on DNA sequencing and metagenomics.

**Figure 4.**
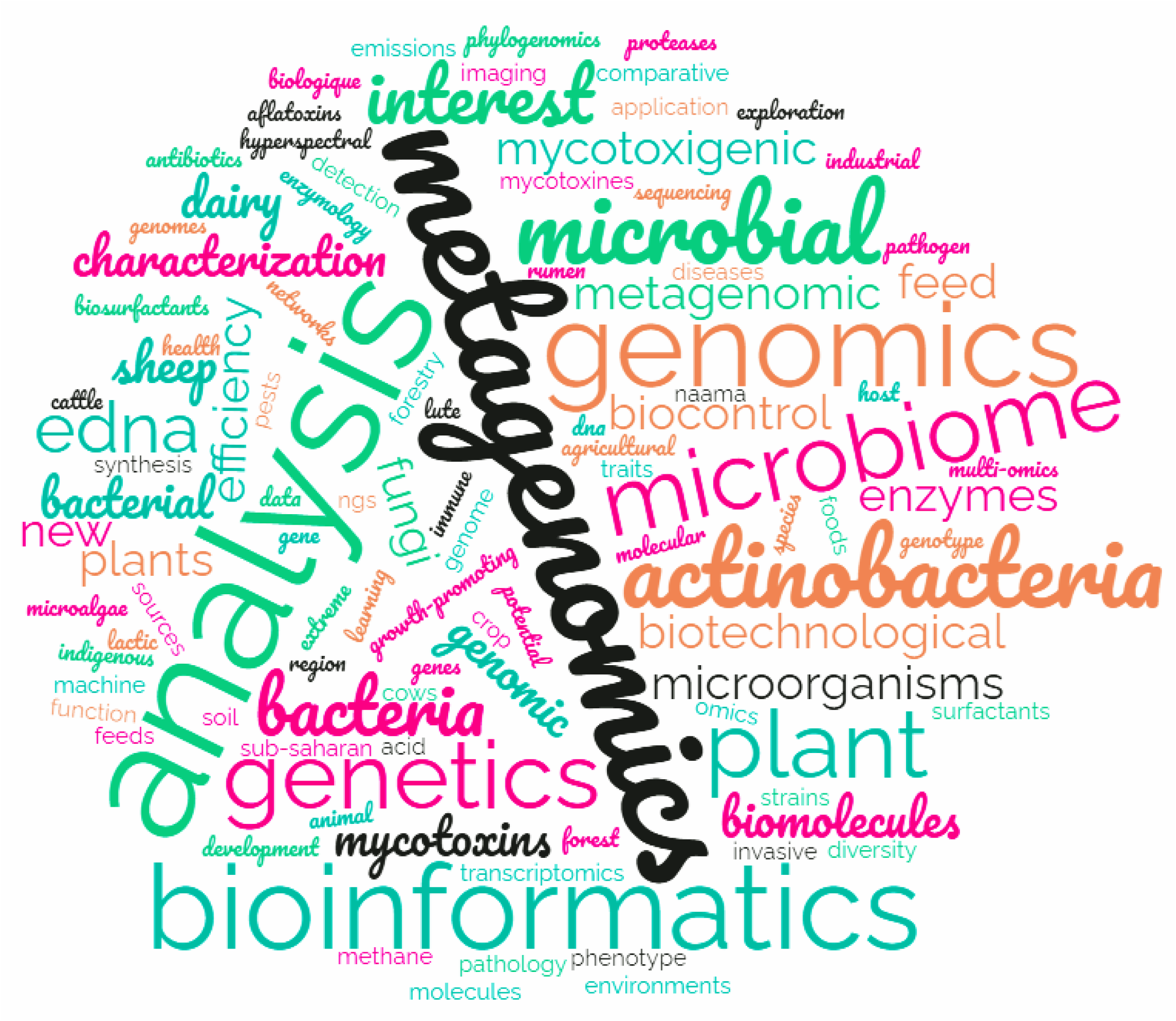
Word cloud of participants’ pre-course stated interests. Visual representation of pre-course interests in specific scientific categories represented using a Word cloud. These data range from the smallest font (least mentioned/interested) to the largest font (most mentioned/interested). Colours show no meaning or impact towards the data and should be ignored.

**Figure 5.**
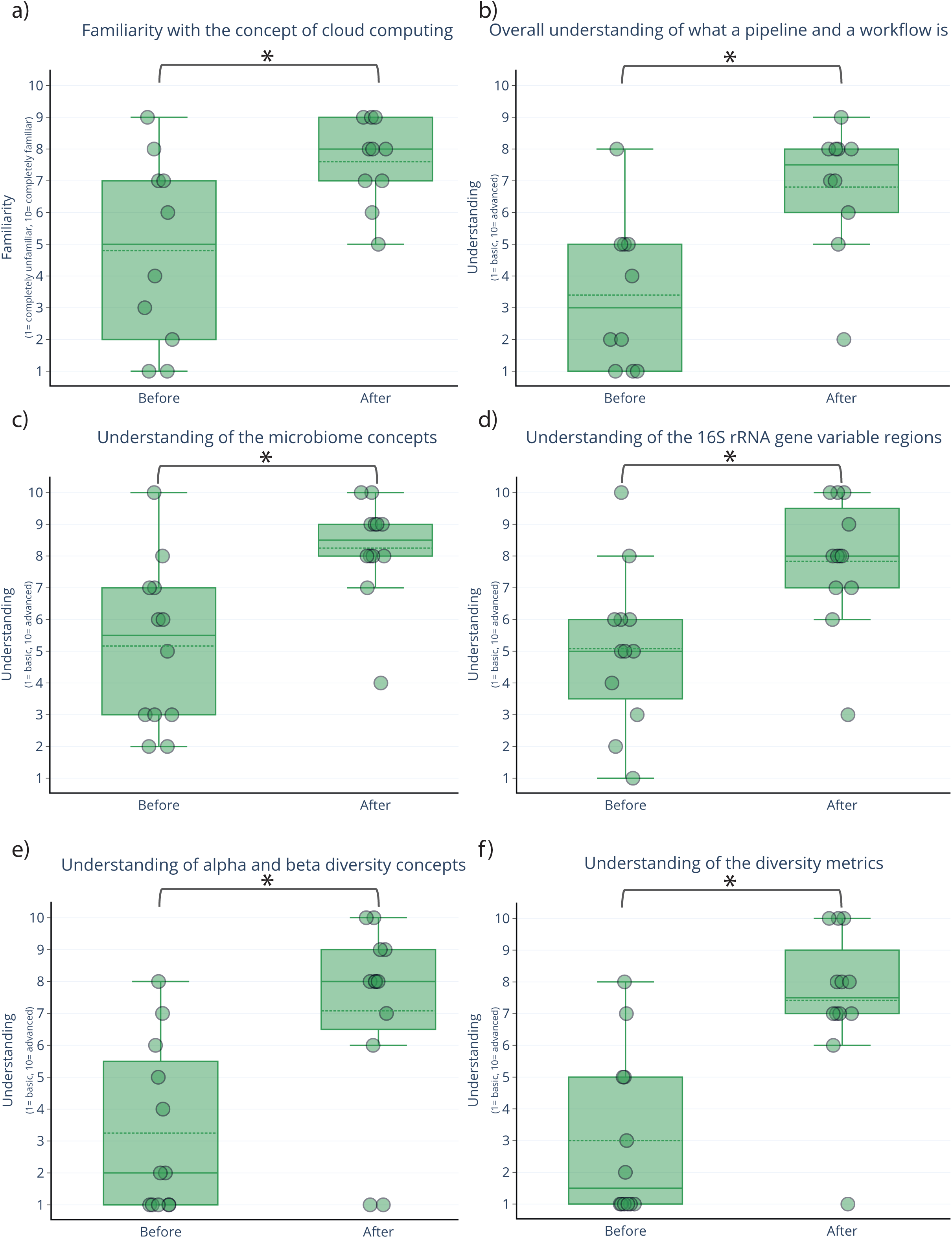
Self-assessments of participants’ confidence in key bioinformatics concepts related to microbiome analysis before and after the course. Box plots showing self-assessment across key concepts in microbiome analysis, before and after attending the event. For each box: upper square line = upper quartile (q3), lower square line = lower quartile (q1), middle square line = median, upper whisker = highest observed value, lower whisker = lowest observed value; dashed middle line = mean. Self-assessment scores range from 1 (basic) to 10 (advanced), with all pre- and post-course comparisons **(b** to **f)** assessed using a permutation test and effect-size estimation using the rank-biserial correlation, except for ‘familiarity with the concept of cloud computing’ evaluation **(a)**, that was assessed for significance using a paired T-test, with effect size determined using Cohen’s d. A significant difference was observed across all comparisons, with a large effect.

Bioinformatics and next-generation sequencing (NGS) are also key areas of interest. Participants are using these technologies for pathogen detection, microbial diversity analysis, and advanced applications like machine learning. The course’s inclusion of cloud-based bioinformatics platforms has catered to both novice and advanced researchers.

Plant and animal health research, such as forest genetics and rumen microbial metagenomics, underscores the need for modules addressing genomics, transcriptomics, and multi-omics approaches. Additionally, many participants are interested in biotechnological applications, including the discovery of new biomolecules in extreme environments, reinforcing the importance of practical bioinformatics tools.

The interest in comparative genomics and phylogenomics further highlights the demand for modules on microbial diversity and genome comparison.

On the other hand, participants expressed a strong likelihood (7.93 ± 2.77, on a scale of 10) of engaging in metagenomics or microbiome-related projects in the next few years (supp. fig. 4). This highlights the relevance of the workshop for their future work, suggesting that the knowledge and skills gained will be directly applicable to their research.

#### 7.4. Self-assessment of confidence and learning progress

##### Cloud computing and workflow construction

The initial familiarity levels with cloud computing was self-assessed as moderate amongst participants (4.8 ± 2.97, on a scale of 10), then improved with an average evaluation at 7.6 ± 1.34 out of 10. A paired t-test revealed a significant improvement in confidence post-course (t = -4.023, p = 0.003), with a large effect size (Cohen’s d = -1.272) (tab. 1). This reflects the course’s effectiveness in providing foundational knowledge on cloud-based solutions. A similar improvement was seen in participants’ understanding of workflows and pipelines, with confidence increasing from 3.4 ± 2.36 before the session to 6.8 ± 2.04 (out of 10) afterward. A permutation test yielded a p-value of 0.047, suggesting a significant improvement in confidence post-intervention. The Rank-Biserial correlation (1.000) indicated a large effect size (tab. 1). This shows that the course’s hands-on sessions were instrumental in helping attendees understand the construction of workflows and pipelines, in a relatively short amount of time.

##### Microbiome-specific learning

Initial understanding of microbiome-related concepts (5.16 ± 2.58, out of 10) increased strongly (8.25 ± 1.60, out of 10) by the end of the course, reflecting the efficacy of both theoretical and practical sessions in improving participants’ comprehension of microbial ecology. A permutation test revealed a significant difference (p = 0.016), with a large effect size (Rank-Biserial correlation = 1.000) (tab. 1). Similarly, familiarity with 16S rRNA gene variable regions improved (from 5.08 ± 2.46, to 7.83 ± 1.99, on a scale of 10) significantly, confirmed by a permutation test (p-value = 0.027), with a large effect size (Rank-Biserial correlation = 1.000) (tab. 1). This jump reflects how well the course helped demystify key areas of microbiome analysis through hands-on learning.

Participants entered the course with minimal knowledge of alpha and beta diversity (3.25 ± 2.63, out of 10), but by the end of the course, their confidence had more than doubled (7.08 ± 3.05, out of 10). A permutation test confirmed a significant difference (p = 0.027), with a large effect size (Rank-Biserial correlation = 1.000) (tab. 1). This indicates that the practical sessions—especially those focused on diversity metrics and visualization—helped demystify complex concepts. Confidence in understanding diversity metrics increased from 3.0 ± 2.59 to 7.41 ± 2.42 (out of 10), with a permutation test indicating a significant difference (p = 0.016), with a large effect size (Rank-Biserial correlation = 1.000) (tab. 1). This further highlights the course’s effectiveness in improving participants’ confidence in understanding microbiome analysis outputs.

#### 7.5. Overall confidence in key concepts

Across all key microbiome analysis concepts, the average confidence level rose from 3.93 ± 1.03 out of 10 before the course to 7.41 ± 0.52 out of 10 after the course (fig. 6). A paired t-test revealed a strong and significant improvement in overall confidence (t = -14.217, p = 0.000), with an exceptionally large effect size (Cohen’s d = -5.374) (tab. 1). The improvement was not only statistically significant but also practically significant, as evidenced by the large effect size. All statistical tests revealed highly significant improvements in participants’ understanding of key scientific concepts following the workshop. These results strongly suggest that the workshop was effective in enhancing participants’ confidence in theoretical and practical knowledge across a range of microbiome-related topics, as well as practical skills such as metadata preparation (a significant improvement of the confidence in preparing a metadata file from 2.83 ± 2.08 to 6.91 ± 1.92 on a scale of 10, with a permutation test p-value = 0.016, and a large effect size according to the rank-biserial correlation p-value = 1.000; supp. fig. 4). The results underscore the educational efficacy of the workshop and its potential for application in similar educational settings.

**Figure 6.**
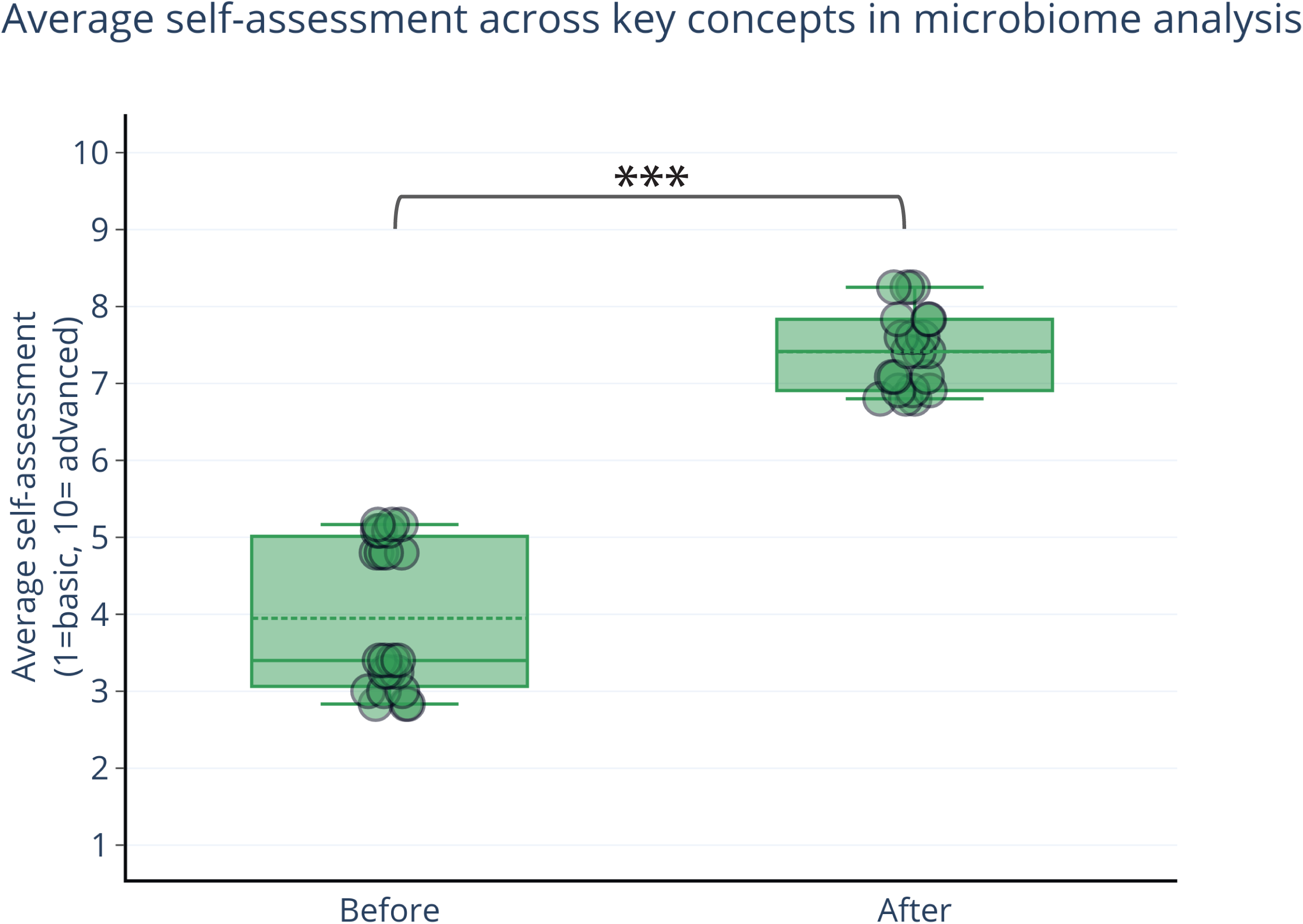
Average self-assessment of participants’ confidence in key bioinformatics concepts related to microbiome analysis before and after the course. Box plots showing average self-assessment across all questioned concepts in microbiome analysis, before and after attending the event. For each box: upper square line = upper quartile (q3), lower square line = lower quartile (q1), middle square line = median, upper whisker = highest observed value, lower whisker = lowest observed value; dashed middle line = mean. Pre- and post-course differences were evaluated using a paired T-test. Effect size was calculated using Cohen’s d. A significant difference was observed across all comparisons, with a large effect.

#### 7.6. Overall acquired knowledge evaluation

To assess the effectiveness of the “Metagenomics Days” event in enhancing participants’ knowledge and skills in metagenomics and bioinformatics, we conducted an anonymous evaluation based on participants’ responses to assessments during the course. Through the post-course survey, these assessments were designed to gauge the understanding and retention of key concepts introduced throughout the lectures and hands-on sessions.

The results (in percentages of total answers) of the assessments (fig. 7) indicated the following:

- **‘Correct’ responses:** 64.87% ± 23.02
- **‘Incorrect’ responses:** 9.35% ± 10.08
- **‘Not sure’ responses:** 25.76% ± 16.05

**Figure 7.**
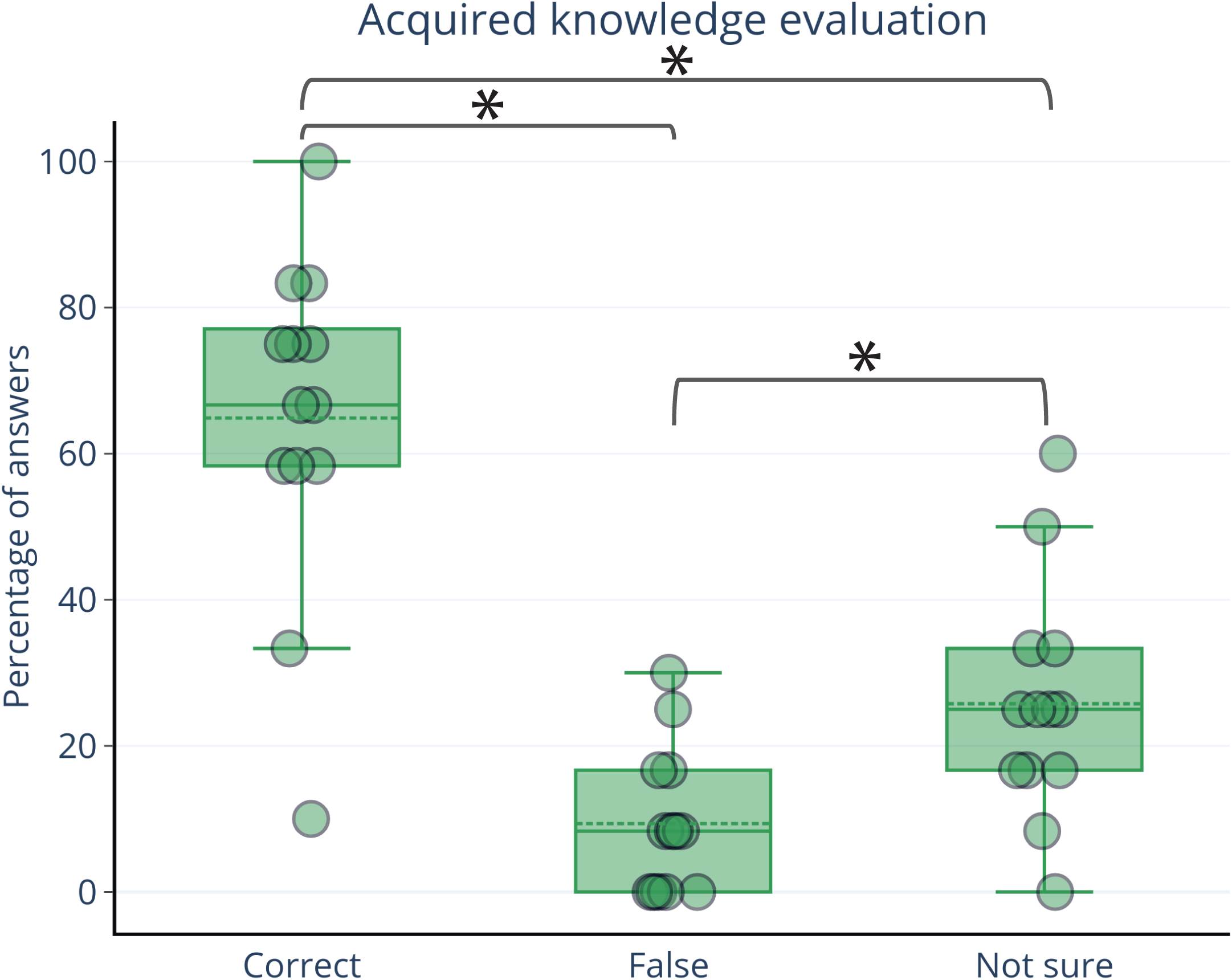
Post-course assessment of participants’ knowledge acquisition. Box plots showing acquired knowledge evaluation of attendees after attending the event. For each box: upper square line = upper quartile (q3), lower square line = lower quartile (q1), middle square line = median, upper whisker = highest observed value, lower whisker = lowest observed value; dashed middle line = mean. Categories include ‘Correct’, ‘False’, and ‘Not Sure’. These categories are displayed as percentage of answers. Friedman test was applied to compare response categories, indicating a significant difference between groups. Wilcoxon signed-rank tests were conducted to assess differences between pairs of answer categories (correct vs. false, correct vs. unsure, false vs. unsure), revealing significant differences across categories, with large effect sizes calculated using the rank-biserial correlation.

A Friedman test indicated a significant difference between groups (χ² = 16.000, p = 0.000). Wilcoxon signed-rank tests revealed significant differences between the correct vs. false (p = 0.0022), correct vs. not sure (p = 0.0242), and false vs. not sure (p = 0.0151) categories, all with large effect sizes (correct vs. false: r = 0.956, correct vs. not sure: r = 0.802, false vs. not sure: r = 0.967) (tab. 1).

These outcomes reflect a significant level of knowledge acquisition among participants, particularly given the complexity of the subject matter and the diversity of participants’ prior experience. The high percentage of correct responses (64.87%) demonstrates that most participants successfully grasped the essential concepts covered in the event, particularly in areas such as sequencing technologies, cloud-based bioinformatics, and microbiome analysis workflows.

The relatively low percentage of incorrect responses (9.35%) suggests that misunderstandings were minimal, likely due to the structured, step-by-step approach of the practical sessions and the continuous availability of instructors for guidance. However, the “Not Sure” responses (25.76%) highlight areas where further clarification and reinforcement of concepts could be beneficial. This may indicate that some topics, particularly those requiring more advanced bioinformatics knowledge (e.g., statistical analysis of microbiome diversity metrics), could benefit from additional time or post-event support, such as follow-up tutorials or online resources.

These results suggest that the course was not only successful in elevating participants’ confidence in understanding of metagenomics workflows and analysis tools, but also in achieving a significant shift in knowledge, with room for improvements in clarifying more complex topics. This feedback will guide future iterations of the event, allowing for targeted improvements in the design and delivery of course content.

Overall, the improvement in confidence and acquired knowledge suggests that the course structure—combining lectures with practical, hands-on sessions, primarily utilizing cloud computing integrative platforms—was highly effective in building participants’ confidence, understanding, and skills.

## VI. Discussion

### Our approach

The approach taken during the “Metagenomics Days” course was distinctive in several key aspects that set it apart from typical educational offerings in metagenomics and bioinformatics. One of the hallmark elements of our training was the adaptation of the QIIME2 Atacama exercise to the EDGE bioinformatics platform. This adaptation was not merely a technical shift; it represented a thoughtful redesign of the learning experience to prioritize user-friendliness. By leveraging EDGE, we provided an intuitive interface that reduced barriers for participants who may have felt overwhelmed by more complex bioinformatics software. This approach allowed new users to engage with metagenomics amplicon sequencing analysis more comfortably and effectively.

We intentionally created an environment that encouraged participant interaction and engagement. The course was designed to facilitate discussions not only during practical sessions but also in the lectures. This interactive learning model fostered a collaborative atmosphere where participants felt free to share their own experiences and challenges. By connecting the course content to their individual research projects, we made the material relevant and applicable, enhancing retention and comprehension.

While many courses, due to shorter time periods, would be compelled to skim the surface of topics in metagenomics, we aimed for a comprehensive understanding, as the four-day program allowed for in-depth exploration. We not only covered foundational principles but also delved into more advanced topics such as quality control, metadata preparation, and the interpretation of complex visualizations like alpha-rarefaction curves. Our course uniquely blended theory with practical application. Each lecture was strategically followed by hands-on sessions where participants could immediately apply what they learned. This immediate application reinforced the concepts and provided a clearer understanding of how to conduct their analyses in real-world scenarios. Recognizing the importance of data visualization in bioinformatics, we dedicated significant time to teaching participants how to interpret various types of charts and graphs that are common within the metagenomics realm. By focusing on the visual representation of data, we aimed to equip participants with the skills to communicate their findings effectively and make informed decisions based on their analyses.

One of the key advantages of using EDGE bioinformatics is its accessible graphical user interface (GUI), which allows users to execute complex bioinformatic pipelines without needing in-depth programming knowledge. The interface supports drag-and-drop functionality, automated workflows, and dynamic visualizations, which makes it ideal for educational settings where the focus is on learning and applying concepts rather than mastering technical intricacies.

Several participants specifically mentioned that they had been hesitant to dive into bioinformatics due to the intimidating nature of command-line interfaces. The streamlined, GUI-driven experience of EDGE not only lowered this barrier but also gave participants the confidence to engage deeply with the biological aspects of their data. They could focus on learning the science of microbial diversity, taxonomic classification, and community analysis without being bogged down by technical hurdles.

Beyond the technical aspects of the course, one of our overarching goals was to foster a sense of community among the participants. Throughout the course, we encouraged collaboration, discussion, and peer support. Participants were given the opportunity to present their research questions and challenges during the sessions, which facilitated a rich exchange of ideas and best practices.

### Challenges

Despite the overall success of the “Metagenomics Days” event, several challenges arose that impacted the smooth execution of the program. Nevertheless, these issues provided valuable insights into areas for improvement in future iterations of the workshop.

Reliable internet access was an essential aspect of the workshop, given the reliance on web-based bioinformatics platforms such as Galaxy and EDGE Bioinformatics. However, inconsistent connectivity proved problematic at times, particularly when participants were working with online tools for sequence analysis and data visualization. This was especially challenging for remote participants who depended entirely on internet access for their engagement.

One limitation of this study is the small sample size of participants, which may affect the generalizability of the findings. Future studies with larger sample sizes are recommended to confirm these results. Additionally, the self-reported nature of the data could introduce bias, and more objective measures of understanding could be considered in future research.

Maintaining flexible coffee break schedules proved difficult due to the variable pacing of different sessions. While some sessions ran longer than expected due to deep discussions or additional hands-on troubleshooting, others finished earlier, resulting in inconsistent break times and participant fatigue during longer segments. The workshop involved a dynamic mix of lectures, practical sessions, and Q&A segments, which made it challenging to adhere strictly to the schedule. For remote participants, this was an even bigger hurdle as it was more difficult to manage their participation across time zones.

Although the course was conducted in English, non-native English speakers—especially remote participants—occasionally struggled with the northern Irish accent of the support staff. This created minor communication challenges that slowed the progress of some discussions.

The design of the course, while comprehensive and in-depth, required significant time investment from both participants and instructors. Certain sections, such as the microbiome statistics sessions, ran longer than anticipated. As a result, some hands-on components had to be condensed to stay on schedule.

The course was primarily delivered by two instructors (Dr. Belaouni and Dr. Stevenson), placing considerable strain on them to handle both lecture delivery and hands-on supervision. The shortage of lecturers made it difficult to provide more individualized attention to participants. Given the shortage of organisational and support staff, the degree of assistance available to remote participants was limited. This occasionally left them without real-time feedback during certain tasks, particularly during more complex hands-on exercises. This was particularly evident when managing computational tasks and software installations, as well as when remote participants experienced technical difficulties.

Space was a limiting factor in the selected physical venue, which was capable of hosting around twenty participants. Fortunately, the ten in-person attendees fit comfortably into the space, facilitating a close-knit atmosphere that fostered active networking and collaboration. Despite the logistical and technical challenges faced, a commendable level of follow-up and engagement was maintained with remote participants.

From a technical point of view, participants using corporate laptops faced significant challenges due to stringent cyber security settings. These restrictions hindered the streamlined sharing of files and datasets, causing delays in certain exercises, particularly those involving the installation and use of third-party software tools. While cyber security restrictions limited the ability to share data via cloud-based systems, the organizers employed a workaround by distributing USB keys. These USB drives were used to share datasets and materials when other methods were unavailable, ensuring that participants could continue the hands-on exercises without significant delays.

### Strength points of the Metagenomics Days event

In addition to the challenges, the Metagenomics Days event demonstrated several key strengths that contributed to its overall success and impact. These strengths played an essential role in ensuring that the course objectives were met while fostering a productive and engaging learning environment.

The use of printed materials, such as manuals, banners, and posters, elevated the professional image of the event. This meticulous attention to detail gave participants the impression of a well-organized and high-quality workshop, reflecting the commitment of the organizing team to excellence.

One of the event’s standout features was the team’s ability to make necessary and astute adjustments during the course. Whether it was adapting to time constraints, managing the hands-on sessions more efficiently, or addressing technical issues as they arose, the flexibility and responsiveness of the organizers allowed the event to run smoothly, even when unforeseen challenges presented themselves.

The cloud computing lecture was lighter than expected, which turned out to be a beneficial adjustment. This allowed participants to engage more easily with the material and stay focused during what could have otherwise been a highly technical and potentially overwhelming session.

Participants were grouped into two-person teams for the practical sessions, which significantly enhanced the pace and effectiveness of the exercises. Working in pairs allowed for peer support, quick troubleshooting, and more dynamic interaction with the course material. This strategy enabled participants to move faster through the steps while gaining a deeper understanding through collaboration.

One of the most enriching aspects of the event was the contribution of participant datasets. Attendees brought in their own RNA-seq datasets, which were used during the quality control (QC) and trimming stages of the hands-on sessions. Although the focus of the workshop was on amplicon sequencing, working with RNA-seq datasets added a layer of complexity and variety, enriching the learning experience. Instructors were careful to highlight the key differences in standards and expectations between RNA-seq and amplicon sequencing libraries, ensuring that participants understood the nuances of both.

The event generated significant interest from the Agri-Food and Biosciences Institute (AFBI) community, particularly among active researchers and scientific staff. By hosting the event in one of the main communal areas of AFBI’s Newforge site, the workshop attracted attention and participation from key members of the institute, further contributing to the interdisciplinary dialogue.

Among the bioinformatics platforms explored during the workshop, Galaxy stood out for its ease of use, particularly in terms of sharing datasets and workflows. While the EDGE bioinformatics platform was also introduced, Galaxy’s user-friendly interface and seamless sharing capabilities allowed participants to engage more effectively with the content.

The success of the event was further bolstered by the support provided by AFBI’s administration. From logistical backing to providing the necessary resources, AFBI played a critical role in the organization and execution of the workshop, ensuring that the event could take place under optimal conditions. Remarkably, the event was run at a very low cost. This was achieved by minimizing expenditures on venue, materials, and technical resources, and by leveraging the institutional support of AFBI. The cost-efficiency of the event made it accessible to a broad range of participants without sacrificing the quality of the content or instruction.

## V. Conclusion, future directions, and suggestions

The “Metagenomics Days” event was a bold step toward democratizing metagenomic amplicon sequencing analysis for both novice and experienced researchers, within AFBI but also amongst external participants. It combined well-structured lectures, hands-on practical sessions, and interactive discussions, which enhanced participants’ learning and engagement. There were some areas for improvement and future directions that could make subsequent events even more successful.

For future hybrid events, we strongly recommend having at least two designated staff members to support remote attendees. This will ensure that their questions are addressed in real-time, technical issues are managed promptly, and they feel fully integrated into the sessions.

In future iterations, it would be essential to have at least one person dedicated solely to handling the technical aspects of the event. This individual would manage audiovisual equipment, ensure smooth streaming for remote participants, and troubleshoot any IT-related problems that arise.

With only two lecturers handling all the sessions, the workload was considerable. For future events, it is recommended to include more lecturers, each specializing in specific aspects of metagenomics, bioinformatics, or microbial ecology. This will not only diversify the content delivered but will also reduce the pressure on individual lecturers, allowing for more focused and detailed discussions on each topic.

To accommodate more participants and expand the reach of the event, we propose the use of two separate venues. One venue can be dedicated to lectures, which can host a larger audience, some of whom may not necessarily participate in the practical sessions. A second venue, designed for computing activities, can then be reserved for hands-on practical work in smaller, more focused groups.

Introducing a paid registration model could enhance interest in the event and bring additional value to participants’ current projects, especially within the context of AFBI. Payment would allow for more professional development opportunities, including better resources and more extensive technical support. At the same time, provisions should be made for reduced fees or full waivers based on merit and financial need, particularly for AFBI staff, students, and researchers with limited funding.

Recording the sessions, especially the lectures, would be highly beneficial for participants who wish to revisit the material or for those unable to attend live. These recordings can also serve as valuable resources for future cohorts or for the institution’s educational repository.

The diversity of responses from participants suggests that the course should be designed with flexibility and tiered learning in mind. It should provide both foundational knowledge for beginners and more advanced modules for those with specific interests in bioinformatics, genomics, and metagenomics. Participants are eager to learn about both the theoretical and practical aspects of these fields, particularly in relation to sequencing technologies, data analysis workflows, and the application of bioinformatics tools.

The success of the “Metagenomics Days” event, particularly through our strategic adaptation of the QIIME2 Atacama exercise to the EDGE bioinformatics platform, highlights the potential of combining powerful, established bioinformatics tools with user-friendly platforms. This approach not only lowers the barriers to entry for newcomers but also enhances the learning experience by making complex workflows more accessible and interactive. Our approach has proven to be both scientifically rigorous and pedagogically effective, enabling participants to gain practical skills in metagenomics that they can immediately apply in their research.

*****

## Supporting information

Announcements

banners

## Acknowledgments

We would like to extend our sincere gratitude to the staff and administrators of the Agri-Food and Biosciences Institute (AFBI) for their invaluable support in facilitating this workshop. Their assistance and dedication ensured the smooth organization and execution of this event. We are also deeply grateful to all participants who attended, engaged actively, and provided feedback, which was essential in refining and enhancing the workshop content. Lastly, we acknowledge the broader metagenomics and microbiome analysis communities for their contributions to open-source software and educational materials, which continue to inspire and empower learning in this rapidly evolving field.

*****

## Supplementary material

**Supplementary material S. 1: Surveys.**

**Supplementary material S. 2: the “Metagenomics Days”’ program**.

**Supplementary material S. 3: Course material.** This material is available through Figshare (https://figshare.com/ndownloader/files/51364694).

**Supplementary figure 1.** Participants’ ability to bring laptops to the course; Pie chart displaying attendees’ ability/willingness to bring their personal laptops to the course. Categories with associated colors include: ‘Yes’ in blue, ‘No’ in red, and ‘May be’ in green.

**Supplementary figure 2**. Familiarity with operating systems; Pie chart displaying results reflecting attendees’ familiarity with different operating systems. ‘Windows’ is shown in blue, ‘Linux’ is shown in orange, and ‘Both’ is shown in green. No participant expressed a familiarity with MacOS.

**Supplementary figure 3.** Assessment of participants’ confidence in building a pipeline in Galaxy using Illumina reads. Box plot showing attendees’ confidence in building a pipeline in Galaxy, using Illumina short reads. For each box: upper square line = upper quartile (q3), lower square line = lower quartile (q1), middle square line = median, upper whisker = highest observed value, lower whisker = lowest observed value; dashed middle line = mean. Confidence values range from 1 (No idea how to build one) to 10 (Could easily build one).

**Supplementary figure 4. Participants’ likelihood of engagement in metagenomics/microbiome projects in the next two years.** Box plot showing likelihood of engagement in a project related to metagenomics/microbiome analysis. For each box: upper square line = upper quartile (q3), lower square line = lower quartile (q1), middle square line = median, upper whisker = highest observed value, lower whisker = lowest observed value; dashed middle line = mean. Likelihood is scored from 1 (very unlikely) to 10 (very likely).

**Supplementary figure 5.** Assessment of participants’ confidence in preparing a metadata file; Box plots showing attendees’ confidence in preparing a metadata file before and after attending the course. For each box: upper square line = upper quartile (q3), lower square line = lower quartile (q1), middle square line = median, upper whisker = highest observed value, lower whisker = lowest observed value; dashed middle line = mean. Confidence values range from 1 (No idea about how to prepare one) to 10 (Could easily prepare one).

---

**Supplementary table 1:** Roles of organizers of the “Metagenomics Days” event.

| *Name/Role* | Dr. Hadj Ahmed<br>Belaouni | Dr. Michael<br>Stevenson | Mr. Stewart Russell | Mr. Andrew McClure |
| --- | --- | --- | --- | --- |
| **Primary role** | Lead organizer | Deputy organizer | Organizer | Organizer |
| **Other roles** | Lecturer, teacher | Lecturer,<br>demonstrator | IT | IT |

