## supplementary material S. 2 for "“The Metagenomics Days”: a simplified workshop on amplicon sequencing analysis with open cloud bioinformatics for eDNA and Microbiomes"

AGRI-FOOD & BIOSCIENCES INSTITUTE

Presents

### The Metagenomics Days

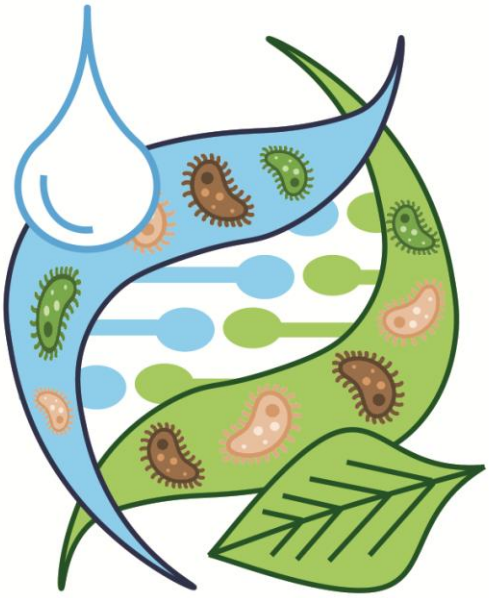

**19-22 February 2024**

Newforge lane, Belfast - NI

### Program of the workshop

#### and guidance on how to reach the training venue

#### Program of the workshop

| | Monday<br>19/02/2024 | Tuesday<br>20/02/2024 | Wednesday<br>21/02/2024 | Thursday<br>22/02/2024 |
|---|---|---|---|---|
| 9:00-10:30 | free | free | free | free |
| 10:30-11:00 | **Registration**<br>& Reception of participants | | | |
| 11:00-12:00 | **DNA sequencing technologies, microbiomes, and e-DNA**<br><br>(first lecture)<br><br>Dr. Belaouni | **Cloud metagenomics: Integrated bioinformatics platforms for metagenomics analysis**<br><br>(second lecture)<br><br>Dr. Belaouni | **Metagenomics amplicon sequencing analysis: an exemplar pipeline**<br><br>(third lecture)<br><br>Dr. Belaouni | **Understanding microbiome metrics**<br><br>(last lecture)<br><br>Dr. Belaouni |
| 12:00-12:30 | **Discussion** | | | |
| 12:30-14:00 | **Lunch Break** | | | |
| 14:00-15:15 | Hands on session: **Understanding sequencing outputs** (part 1)<br><br>Dr. Belaouni & Dr. Stevenson | Hands on session: **Navigating cloud bioinformatics resources** (part 1)<br><br>Dr. Belaouni & Dr. Stevenson | Hands on session: **Amplicon sequencing analysis using cloud bioinformatics** (part 1)<br><br>Dr. Belaouni & Dr. Stevenson | Hands on session: **Exploring QIIME2 outputs** (part 1)<br><br>Dr. Belaouni & Dr. Stevenson |
| 15:15-15:30 | **Coffee & Tea break** | | | |
| 15:30-16:30 | Hands on session: **Understanding sequencing outputs** (part 2)<br><br>Dr. Belaouni & Dr. Stevenson | Hands on session : **Navigating cloud bioinformatics resources** (part 2)<br><br>Dr. Belaouni & Dr. Stevenson | Hands on session: **Amplicon sequencing analysis using cloud bioinformatics** (part 2)<br><br>Dr. Belaouni & Dr. Stevenson | **Exploring QIIME2 outputs** (part 2)<br><br>Dr. Belaouni & Dr. Stevenson |
| 16:30-17:00 | **Discussion** | | | |
| 17:00-17:10 | **Group pictures** | | | **Closure of the event + Group picture (last Day)** |

#### Location of the venue

##### Address

18a, Newforge Lane, Belfast, Co. Antrim, Belfast

BT9 5PX

##### How to reach it?

https://maps.app.goo.gl/iKBGWRknYpLEGb1VA

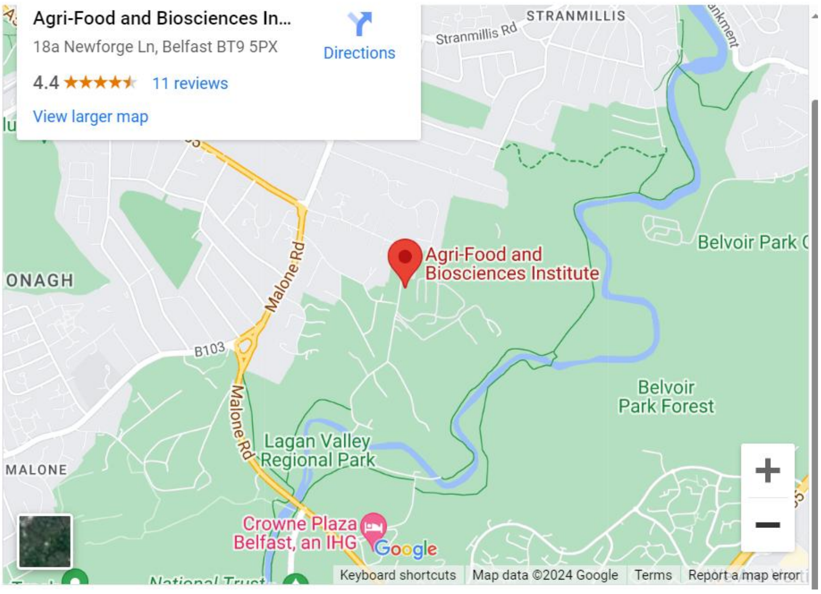


##### By Bus (from Donegall Square, City Center):

https://maps.app.goo.gl/FcKrftuyw8sTpXF69

##### On foot (from Donegall Square, City Centre):

https://maps.app.goo.gl/MmE5S6gL7GSoTQtx5

##### By Car (from Donegall Square, City Centre):

https://maps.app.goo.gl/jRPc4amrtstJjc4C7

#### AFBI’s Newforge site map

Via visitors’ entry, once in the premises, go to the Reception (Faculty Building), and ask about the Foyer Conference Room.

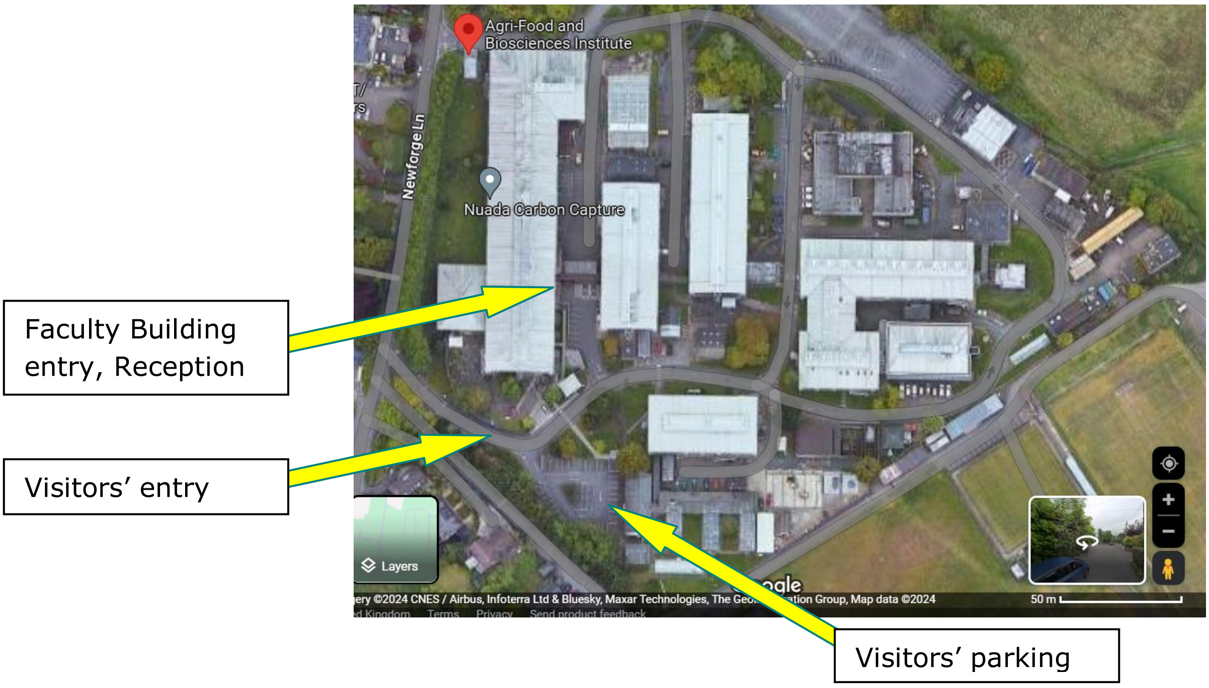


#### Have any inquiry?

Don’t hesitate to reach out:

 (Dr. H.A. Belaouni)

 (Dr. M. Stevenson)


### Welcome to the Metagenomics Days!

The Metagenomics Days
19-22 February 2024
Newforge lane, Belfast - NI
