## Supplementary material for "“The Metagenomics Days”: a simplified workshop on amplicon sequencing analysis with open cloud bioinformatics for eDNA and Microbiomes": Announcements

AGRI-FOOD & BIOSCIENCES INSTITUTE

presents

### The Metagenomics Days

#### an introduction to amplicon sequencing analysis (2024)

19 – 22 February 2024 | Belfast, UK

##### TOPICS

1. DNA sequencing technologies, microbiomes, and e-DNA
2. Cloud metagenomics: Integrated bioinformatics platforms for metagenomics analysis
3. Metagenomics amplicon sequencing analysis: an exemplar pipeline
4. Understanding microbiome metrics

##### ORGANIZERS

Dr. Hadj Ahmed Belaouni
AFBI, UK

Dr. Michael Stevenson
AFBI, UK

**Very Limited Spots - Secure Your Seat Today!**

##### REGISTRATION

Registration deadline
31 January 2024

IMPORTANT DATES

- 17/01/2024: Announcement and 1st Call.
- 24/01/2024: 2nd Call.
- 31/01/2024: Registration deadline.
- 19/02/2024: Workshop commencement.
- 24/02/2024: Workshop conclusion.

##### CONTACT

Hadj Ahmed BELAOUNI


Registration link http://bit.ly/metagenomics-afbi
