## Supplementary material for "“The Metagenomics Days”: a simplified workshop on amplicon sequencing analysis with open cloud bioinformatics for eDNA and Microbiomes": banners

afbi AGRI-FOOD & BIOSCIENCES INSTITUTE presents:

### The Metagenomics Days

#### An introduction to amplicon sequencing analysis (2024)

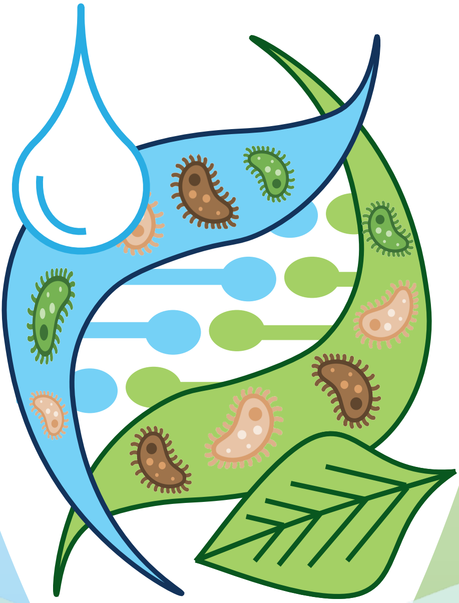


ORGANIZERS

Dr. Hadj Ahmed Belaouni
AFBI, UK

Dr. Michael Stevenson
AFBI, UK


19 – 22 February 2024
Belfast, UK

TOPICS

1. DNA sequencing technologies, microbiomes, and e-DNA
2. Cloud metagenomics: Integrated bioinformatics platforms for metagenomics analysis
3. Metagenomics amplicon sequencing analysis: an exemplar pipeline
4. Understanding microbiome metrics
