## Supplementary figures and images for "“The Metagenomics Days”: a simplified workshop on amplicon sequencing analysis with open cloud bioinformatics for eDNA and Microbiomes"

### supp. fig. 1

#### Would you be able to bring your laptop?

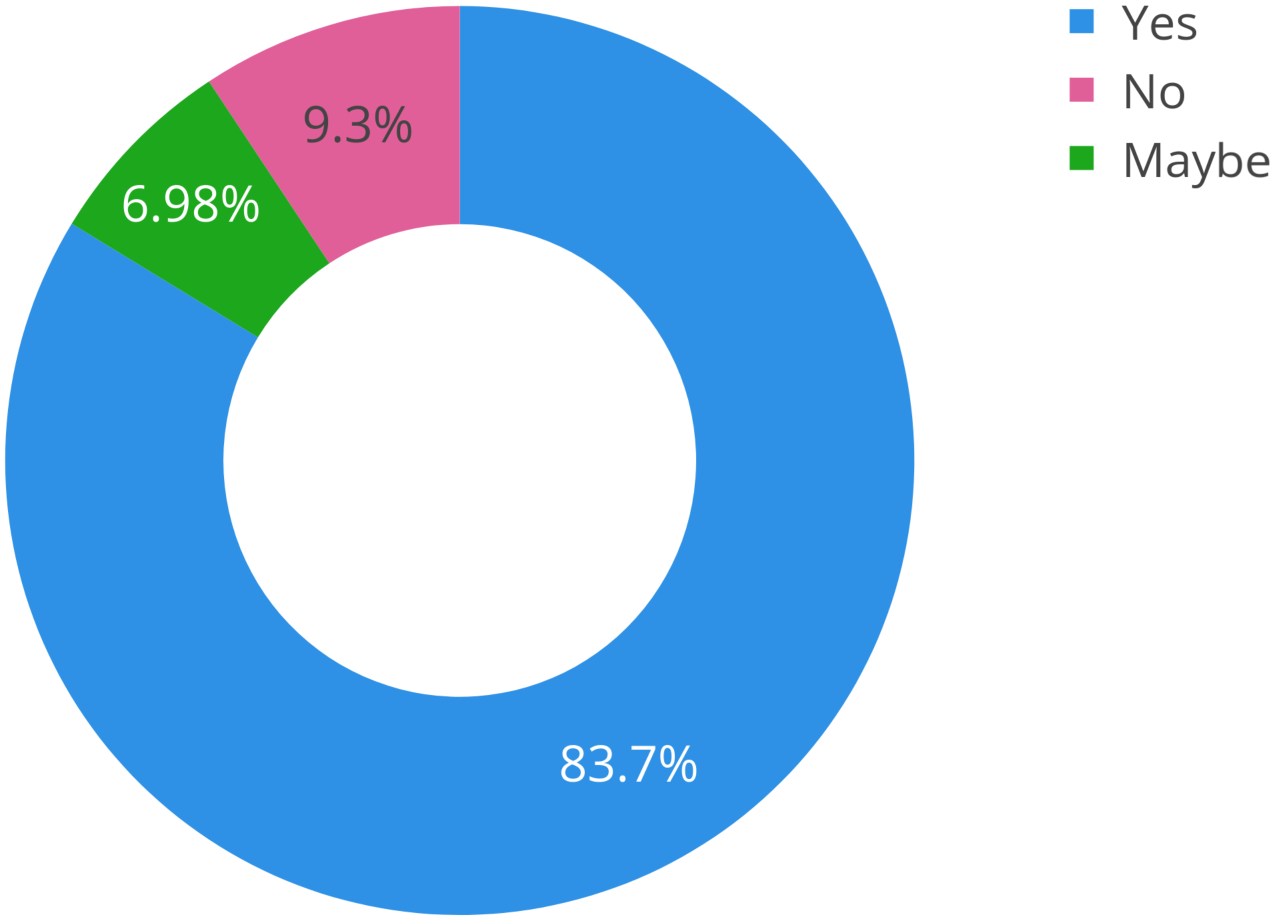

### supp. fig. 2

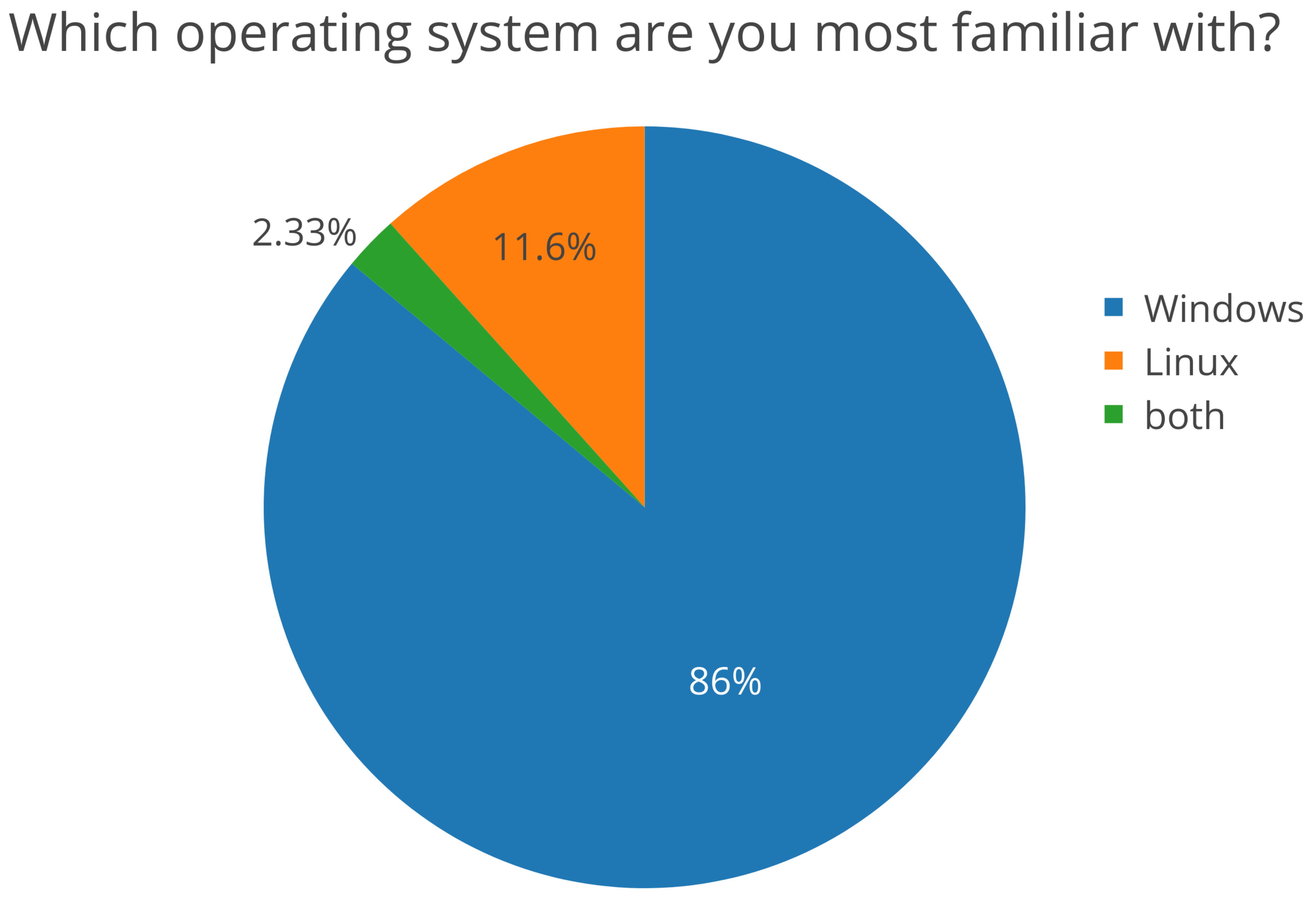

### supp. fig. 3

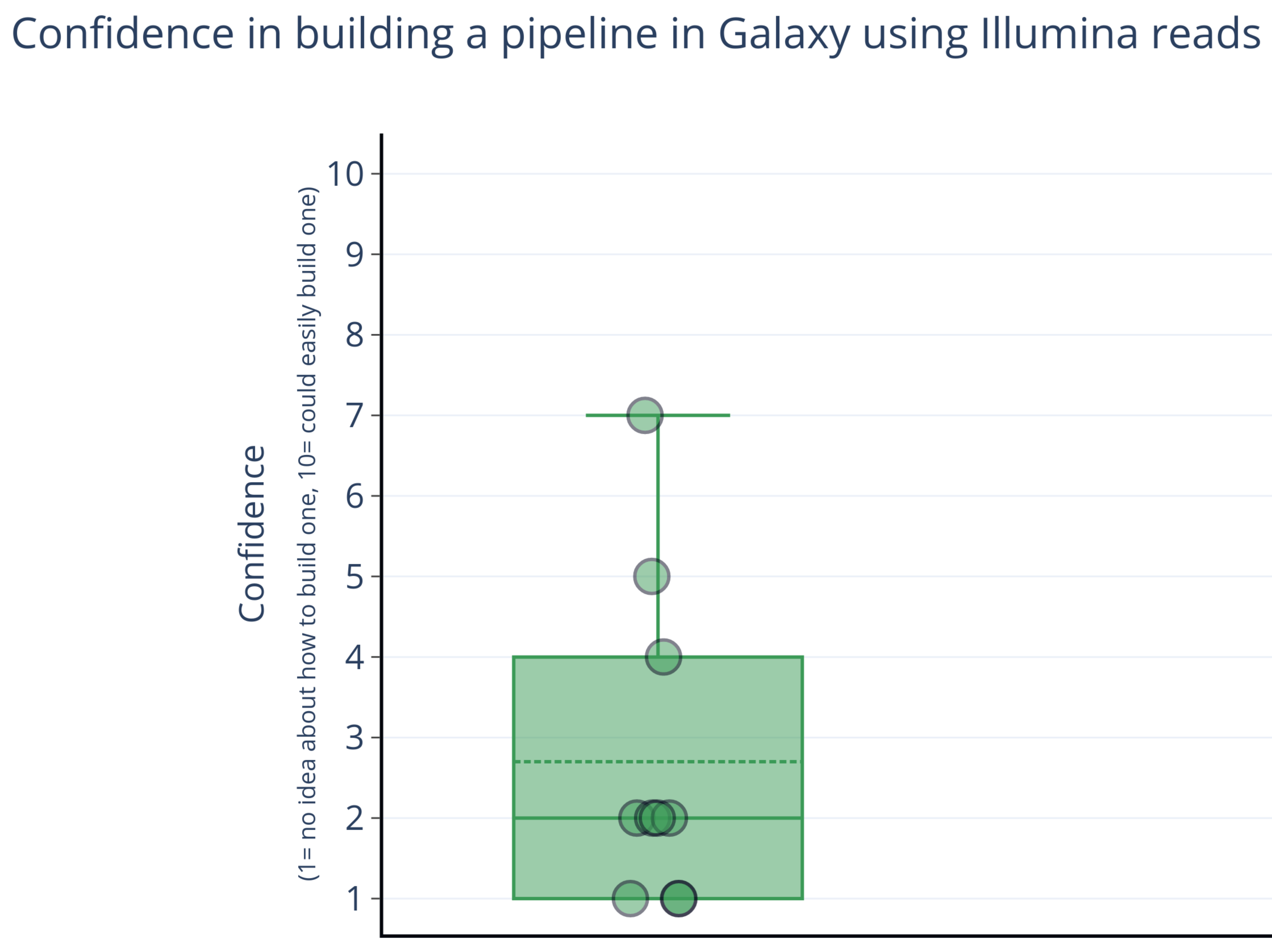

### supp. fig. 4

Likelyhood of engagement in a project related to metagenomics/microbiome analyses

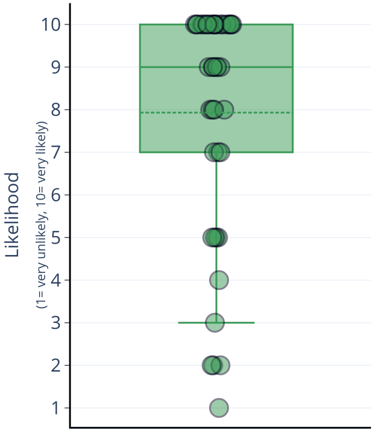

### supp. fig. 5

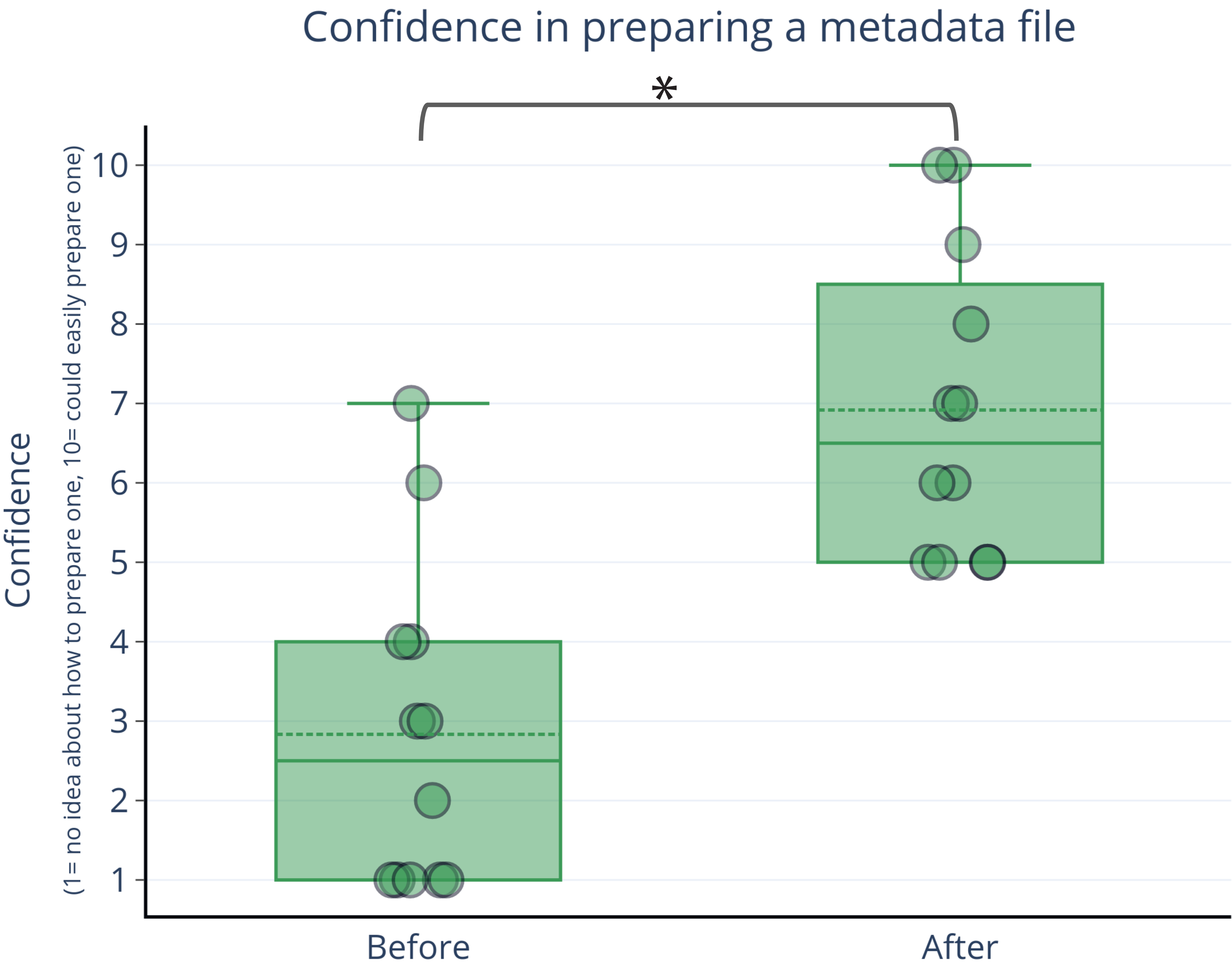
