## supplementary material S. 1 for "“The Metagenomics Days”: a simplified workshop on amplicon sequencing analysis with open cloud bioinformatics for eDNA and Microbiomes"

**Supplementary material S. 1: Surveys**

**I. Pre-course survey**

1. Country (open choices)
2. Current affiliation (open choices)
3. Backgrounds (open choices)
4. Current research interests (open choices)
5. Which one of the four topics is the most important for you?

(Cloud metagenomics and web-based bioinformatics; Metagenomics amplicon sequencing analysis; Microbiome metrics; DNA sequencing technologies, microbiomes, and e-DNA; all of them)

6. What are the subjects you would like to be covered during this event? (open choices)
7. Would you be able to bring your laptop? (Yes/No)
8. Which operating system are you most familiar with? (open choices)
9. What is your primary motivation for taking this course? (open choices)
10. Do you have any specific needs to enhance your experience at this event? (open choices)
11. What are your suggestions for this event? (open choices)
12. Name the bioinformatics resources dedicated to microorganisms that you already have heard of, prior to the course. (Choices: Galaxy; Blast software; R ; Megan; Mega; RDP; NCBI databases; Qiime2; Phyloseq; EDGE; Morpheus; Kbase; None)
13. Which of the following cloud-based tools had you used, prior to this course? (Choices: MEGA; EDGE Bioinformatics ; Kbase; Galaxy; QIIME2; BV-BRC; All of the above; None of the above)
14. Had you ever used a cloud based bioinformatics solution before the course? (No/Yes)
15. Did you know any data repository/database for microbiome studies before the course? (No/Yes)
16. Before this course, were you aware of QIIME2 and what is it used for? (No/Yes)
17. On a scale from 1 (basic) to 10 (advanced), how do you evaluate your understanding of Sanger sequencing technology?
18. On a scale from 1 (basic) to 10 (advanced), how do you evaluate your understanding of Next Generation Sequencing (NGS) technologies?
19. On a scale from 1 (novice) to 10 (expert), how do you evaluate your informatics skills?
20. On a scale from 1 (novice) to 10 (expert), how do you evaluate your BIOinformatics skills?
21. Do you have previous experience with NGS? (No/Yes)
22. Do you have past experience with genomics, metagenomics, or transcriptomics? (No/Yes)
23. On a scale of 1 (very unlikely) to 10 (highly probable), how probable is it that you will be engaged in a project related to metagenomics/microbiome analyses in the next couple of years?
24. How confident were you in building a pipeline in Galaxy using Illumina reads before yesterday's tutorial?
25. Any other comment? (open choices)

**II. Post-course survey**

**a. Confidence/understanding self-assessment**

1. How familiar were you with the concept of cloud computing before day 1 of the course?
   (1= completely unfamiliar, 10= completely familiar)
2. How familiar are you with the concept of cloud computing now?
   (1= completely unfamiliar, 10= completely familiar)
3. How confident were you in building a pipeline in Galaxy using Illumina reads before yesterday's tutorial?
   (1= no idea about how to build one, 10= could easily build one)
4. How would you rate your overall understanding of what a pipeline and a workflow is, prior to yesterday's sessions?
   (1= basic, 10= advanced)
5. How would you rate your overall understanding of what a pipeline and a workflow is now?
   (1= basic, 10= advanced)
6. How well did you understand the microbiome before the workshop?
   (1= basic, 10= advanced)
7. How well do you understand the microbiome now?
   (1= basic, 10= advanced)
8. How well did understand the 16S rRNA gene variable regions before the workshop?
   (1= basic, 10= advanced)
9. How well do you understand the 16S rRNA gene variable regions now you have been to the workshop? (1= basic, 10= advanced)
10. How confident were you about preparing a metadata file before day 3’s hands-on session?
    (no idea about how to prepare one, 10= could easily prepare one)
11. How confident are you about preparing a metadata file now you have attended the workshop?
    (no idea about how to prepare one, 10= could easily prepare one)
12. How confidently did you understand alpha and beta diversity prior to the workshop?
    (1= basic, 10= advanced)
13. How confidently do you understand alpha and beta diversity now you have attended the workshop?
    (1= basic, 10= advanced)
14. How well did you understand the diversity metrics before the workshop?
    (1= basic, 10= advanced)
15. How well do you understand the diversity metrics now you have attended the workshop?
    (1= basic, 10= advanced)

**b. Knowledge-based queries**

*(The answers were evaluated, then labeled as: ‘Correct’, ‘False’, or ‘Not sure’ depending on the participant’s answer)*

1. What is the difference between Qiime2 and Kraken?
2. What is the difference between amplicon metagenomics and shotgun metagenomics ?
3. Why do we use the 16SrRNA for bacterial communities profiling studies?
4. Are the variable regions of the 16S all equally informative?
5. Which combination of 16S variable regions is the most widely used for microbiome analysis?
   (choices: 16S rRNA variable regions)
6. Can we use artifacts as inputs for Qiime2 methods?
7. Can we use visualizations as inputs for Qiime2 methods?
8. Which of the following captures more variations within 16S rRNA sequences?
   (choices: clustering methods)
9. What is the central output of the microbiome analysis in Qiime2?
10. Can we compare the outputs of a study that used denoising with a study that used OTU clustering?
11. Name one alpha diversity metric that is phylogeny aware.
12. Name one beta diversity metric that is qualitative.
13. Name one alpha diversity metric that is quantitative.

*****
